# Unique and shared functions of the Rad9–Hus1–Rad1 and Mre11–Rad50–Nbs1 complexes in ATR checkpoint activation and long-range DNA end resection in *Xenopus* egg extracts

**DOI:** 10.1101/2023.08.10.549595

**Authors:** Kensuke Tatsukawa, Reihi Sakamoto, Yoshitaka Kawasoe, Yumiko Kubota, Toshiki Tsurimoto, Tatsuro S. Takahashi, Eiji Ohashi

**Affiliations:** Graduate School of Systems Life Sciences, Kyushu University, 744 Motooka, Nishi-ku, Fukuoka 819-0395, Japan; Faculty of Science, Kyushu University, 744 Motooka, Nishi-ku, Fukuoka 819-0395, Japan; Graduate School of Science, Osaka University, 1-1 Machikaneyama-cho, Toyonaka, Osaka 560-0043, Japan; Nagahama Institute of Bio-Science and Technology, 1266 Tamura-cho, Nagahama, Shiga 526-0829, Japan

## Abstract

Sensing and processing of DNA double-strand breaks (DSBs) are vital to genome stability. DSBs are primarily detected by the ATM checkpoint pathway, where the Mre11–Rad50–Nbs1 (MRN) complex serves as the DSB sensor. Subsequent DSB end resection promotes the transition from the ATM to the ATR checkpoint pathway, where replication protein A, MRN, and the Rad9–Hus1–Rad1 (9–1–1) checkpoint clamp serve as the DNA structure sensors. 9–1–1 and MRN recruit Topbp1, a critical checkpoint mediator that activates the ATR kinase. However, how multiple sensors contribute to regulating end resection and checkpoint activation remains ambiguous. Using DNA substrates that mimic extensively resected DSBs, we show here that MRN and 9–1–1 redundantly stimulate Dna2-dependent long-range end resection and ATR activation in *Xenopus* egg extracts. MRN serves as the loading platform for Dna2, ATM, and Topbp1. In contrast, 9–1–1 is dispensable for bulk Dna2 loading, and Topbp1 loading is interdependent with 9–1–1 in this pathway. ATR facilitates Mre11 phosphorylation and ATM dissociation. Our results delineate the molecular mechanism of and interplay between two redundant pathways that stimulate ATR checkpoint activation and long-range DSB end resection in vertebrates.

## INTRODUCTION

DNA double-strand breaks (DSBs) are one of the most deleterious types of DNA lesions that cause the fragmentation of chromosomes. Since DSB repair must restore the continuity of broken DNA termini, the repair process requires DSB processing, such as the exposure of single-stranded DNA (ssDNA) regions [1, 2]. Therefore, the DSB sensing machinery must respond to different forms of unprocessed and processed DSBs [3-6]. Defects in the sensing and processing machinery of DSBs give rise to inefficient and inaccurate repair of DSBs. Mutations in these reactions underlie many human genetic disorders and cancers that exhibit genetic instability.

DSBs are repaired mostly through two major repair pathways, non-homologous end joining (NHEJ), which ligate the broken DSB ends with minimal processing, and homology-directed repair (HDR), which uses homologous sequences to restore genetic information [1, 7]. Since HDR requires 3′-terminated ssDNA for homology search, this process involves nucleolytic degradation of the 5′-terminated strand at DSBs. Unprocessed DSBs are first bound by the Ku heterodimer, which serves as the platform for the assembly of the NHEJ machinery [7]. During this step, DSBs are also recognized by Mre11–Rad50–Nbs1 (MRN), a multifunctional complex whose Mre11 subunit has endo- and 3′→5′ exonuclease activities [8, 9]. MRN, together with its activator CtIP, introduces nicks on the 5′-terminated strand and degrades this strand from the nicks by its 3′→5′ exonuclease activity. This “short-range” resection is followed by processive “long-range” resection mediated by two redundant pathways, a combination of DNA helicase/nuclease Dna2 and RecQ family 3′→5′ helicases (BLM and WRN in vertebrates and Sgs1 in yeast) and 5′→3′ exonuclease Exo1, respectively, in the 5′ to 3′ direction [2, 10-15].

The processing of DSBs leads to the transition of the DSB sensing pathways [16, 17]. Minimally processed DSBs are monitored by the ataxia telangiectasia-mutated (ATM) checkpoint pathway, which relies on the ATM serine/threonine-protein kinase [3, 4, 6]. Another related protein kinase, ataxia telangiectasia and Rad3-related (ATR), governs the ATR checkpoint pathway that responds to the exposure of ssDNA [4, 5]. ATM and ATR phosphorylate hundreds of downstream substrates to regulate DNA repair, cell cycle progression, apoptosis, and other cellular reactions. MRN activates ATM mainly through the interaction of Nbs1 with ATM [8, 9]. DSB end resection exposes 3′-terminated ssDNA and generates 5′-terminated primer/template junctions (5′-P/T junctions), promoting the transition from the ATM to the ATR signaling pathway [16, 17].

Importantly, ATR activation depends not only on ssDNA exposure but also on other DNA-structure-specific sensors and a mediator protein that bridges DNA-structure sensors and ATR. ATR forms a complex with ATR interacting protein (ATRIP), which binds to the major eukaryotic ssDNA binding protein, replication protein A (RPA), and functions as an ssDNA sensor [18, 19]. The checkpoint clamp complex, Rad9–Hus1– Rad1 (9–1–1), a ring-shaped heterotrimer that structurally resembles the replication clamp proliferating cell nuclear antigen (PCNA), is loaded by the Rad17-Replication-factor-C-like complex (Rad17-RLC) onto 5′-P/T junctions and serves as the 5′-P/T junction sensor [20-24]. These two sensors are interlinked by Topbp1, a critical mediator protein that carries an ATR-activation domain (AAD) and nine BRCT motifs [25-27]. Some of the BRCT motifs bind phosphopeptides to mediate the phosphorylation-induced multivalent protein-protein interaction [28, 29], which can lead to the formation of protein condensate and activates the ATR checkpoint pathway [30]. The Rad9 subunit of 9–1–1 carries a largely disordered C-terminal tail, which receives phosphorylation on multiple sites [31-33]. Phosphorylation of the specific serines in the C-terminal tail stimulates its interaction with Topbp1 and promotes ATR activation [32, 34-37].

There are significant functional overlaps between DSB-end resection and ATR-activation pathways. In addition to 9–1–1, MRN also activates ATR by recruiting Topbp1 [38-43]. Studies in *Xenopus* egg extracts showed that, in response to poly(dA)_70_–(dT)_70_ synthetic oligonucleotides and restriction enzyme-mediated DSBs, MRN recruits both ATM and Topbp1 to DNA substrates, stimulates phosphorylation of Topbp1 on serine 1131 by ATM, and enhance Topbp1-mediated ATR activation [38, 42, 44]. Similar MRN-dependent Topbp1 recruitment and ATR activation occur on circular ssDNA carrying 5′-P/T junctions in *Xenopus* egg extracts, although 9–1–1 is required for ATR activation as well in this case [39]. Furthermore, MRN facilitates ATR activation in aphidicolin-treated *Xenopus* egg extracts [40] and camptothecin-treated human cells [41]. MRN also facilitates long-range resection by stimulating both the EXO1 and BLM-DNA2 pathways [13, 15, 45, 46]. Likewise, *Xenopus* MRN stimulates the activity of Exo1 and Wrn-Dna2 [47, 48]. Therefore, MRN is involved in ATM activation, short-range and long-range DSB end resection, and ATR activation. Similarly, the checkpoint clamp 9–1–1 regulates DSB end resection [49-52]. In yeast, 9–1–1 restricts the MRN activity to negatively regulate short-range resection [52]. Furthermore, yeast 9–1–1 promotes end-resection by Exo1 and Dna2-Sgs1 at uncapped telomeres [49, 51]. Consistently, purified human 9–1–1 stimulates the enzymatic activity of both DNA2 and EXO1 [49], and 9–1–1 facilitates DSB-end resection in human cells [50]. Therefore, the checkpoint sensors regulate both the activation of checkpoint kinases and DSB end-resection pathways. However, it remains ambiguous how the interplay between multiple checkpoint signaling factors and DSB end resection pathways is regulated and how they are integrated to facilitate robust and efficient sensing and processing of DSBs.

To understand the contribution of 9–1–1 and MRN to DSB sensing and processing pathways, we combined unique DNA substrates that mimic extensively resected DSB termini and the nucleoplasmic extract of *Xenopus* eggs (NPE) that recapitulates many DNA transactions, including DSB end resection and checkpoint activation [53]. We show here that both MRN and 9–1–1 facilitate long-range end resection and ATR activation in NPE. These two pathways function in parallel in response to DSBs but not on circular DNA substrates. Both complexes enhance the loading of ATR and Topbp1 onto DNA. The MRN-dependent ATR activation pathway relies on ATM, while the previously identified serine 1131 phosphorylation site was required for ATR activation only partially. Our data describe molecular mechanisms of how checkpoint sensors regulate both ATR activation and DSB long-range end resection and clarify the unique and shared regulation of these pathways.

## MATERIALS AND METHODS

### *Xenopus* egg extracts

*Xenopus laevis* was purchased from Kato-S-Science (Chiba, Japan) and handled according to the animal experiment regulations at Kyushu University. Preparation of NPE was carried out essentially as described previously [54].

### Cloning and Plasmids

Symbols of *Xenopus* genes and proteins conformed to the nomenclature guidelines of Xenbase (https://www.xenbase.org/entry/static/gene/geneNomenclature.jsp). The *Xenopus laevis rad9a* gene was obtained from the DNASU plasmid repository (#XlCD00712232). *Xenopus laevis hus1, rad1, rad50, nbs1, atm, atr, topbp1, dna2,* and *exo1* genes were amplified from *Xenopus* egg cDNA by PCR using primers listed in Supplementary Table S1. Fusion of the FLAG tag and the TEV protease site and the construction of mutant genes were also carried out by PCR using primers listed in Supplementary Table S1. All the original and modified genes were first cloned into pDONR201 (Thermo Fisher Scientific, MA, USA) by either the Gateway BP reaction or restriction-enzyme-based cloning and then transferred into pET-HSD [54], an in-house Gateway destination vector carrying a T7 promoter and an N-terminal His_6_-tag, for protein expression in *E. coli* and BacuroDirect C-term Linear DNA (Thermo Fisher Scientific, #12562019) for protein expression in insect cells by the Gateway LR reaction. For Topbp1 expression in *E. coli*, the wild-type and the S1131A mutant *topbp1* genes were inserted into the pET28a(+) vector (Merck, Hessen, Germany, #69864) together with a 5′ FLAG-tag sequence.

The expression plasmid for GST-S-LacI was constructed as follows: The *E. coli lacI* gene was amplified from pGEX-6p-3 (Cytiva, MA, USA, #28954651) by PCR using primers eo1190 and eo1193 (Supplementary Table S1), digested with BglII and XhoI, and inserted between the BamHI and XhoI sites in pGEX-6p-3. The resulting GST-S-LacI fusion sequence was then amplified by PCR using primers eo1211 and eo1212 (Supplementary Table S1) and inserted between the NdeI and HindIII sites in the pCold II vector (Takara Bio, Shiga, Japan, #3362) by the In-Fusion reaction, resulting in pColdII-GST-S-LacI.

The plasmid for DNA pull-down substrate preparation (pR3k-lacO4) was constructed as follows: A DNA fragment carrying two BsaI sites and four tandem lacO sites was made by annealing and ligating 5′-phosphorylated oligonucleotides tt2322, tt2323, tt2324, tt2325, tt2326, and tt2327 (Supplementary Table S1). The resulting fragment was then inserted between the HindIII and EcoRI sites in modified pUC119 whose BsaI site had been mutated by overlap-extension PCR using primer pairs tt2318 and tt2321, and tt2319 and tt2320 (Supplementary Table S1).

### Preparation of DNA substrates for checkpoint activation and DNA pull-down assays

DNA substrates for checkpoint activation in NPE were prepared as follows: Primed circular ssDNA was prepared by annealing an 80-mer oligonucleotide (eo1184, Supplementary Table S1, for Figure 1E) or a 38-mer oligonucleotide (tt3253, Supplementary Table S1, for Figure 1B and 1C) onto 3-kb circular ssDNA (pTT336). Primed linear ssDNA was prepared by annealing a 38-mer oligonucleotide (tt3253, Supplementary Table S1) onto linearized pTT336 ssDNA, which was made by annealing a 35-mer oligonucleotide (tt1939, Supplementary Table S1) onto pTT336 ssDNA, digesting it with XhoI, and purifying linearized ssDNA after heat-denaturing the annealed oligonucleotide. Linear dsDNA was prepared by digesting 3-kb and 6-kb plasmids (pTT336 for Figures 1B and 1C, and pTT376 for Figures 1E, 2–4, and 6, and Supplementary Figures S1–4 and S6) with XhoI and ApaI, respectively. To append 3′-ssDNA overhangs, 0.1 µM of the 6-kb fragment was incubated with 1 unit/µL terminal deoxynucleotidyl transferase (TdT, New England Biolabs, MA, USA, #M0315S) and 0.5 mM deoxyadenosine triphosphate (dATP) at 37°C for 30 min. All the DNA substrates were purified by phenol/chloroform extraction and ethanol precipitation and resuspended in TE buffer (10 mM Tris-HCl pH 7.4, 1 mM ethylenediaminetetraacetic acid [EDTA]) at 200 ng/µL. The poly(dA)_70_–(dT)_70_ substrate was prepared by annealing poly(dA)_70_ and poly(dT)_70_ oligonucleotides in TEN buffer (10 mM Tris-HCl pH 7.4, 1 mM EDTA, 50 mM NaCl) at 200 ng/µL. The sequences of pTT336 and pTT376 are available upon request.

**Figure 1.**
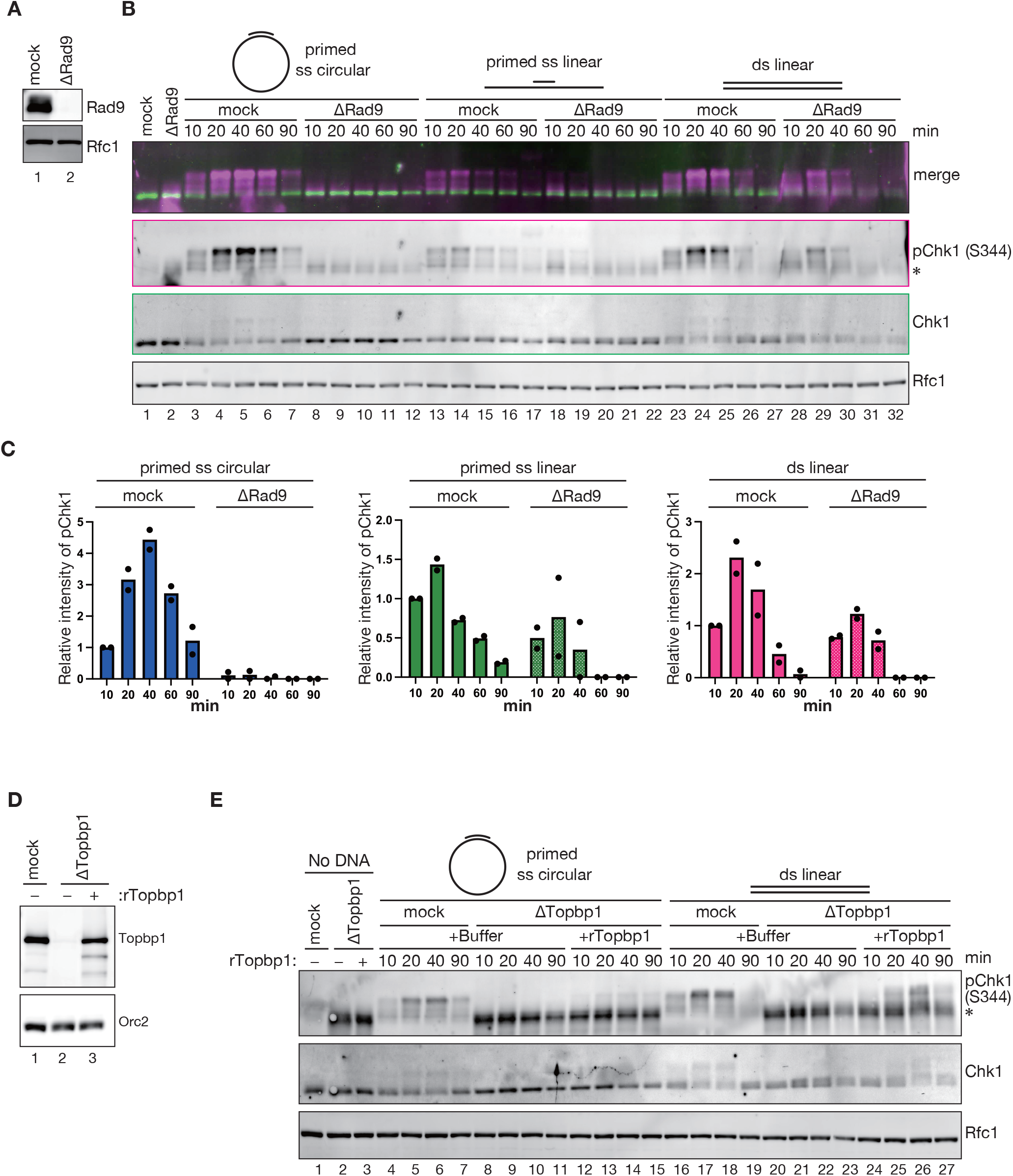
Linear DNA activates ATR through a 9–1–1-independent pathway in Xenopus egg extracts. **(A)** 0.2 µL each of mock-treated (lane 1) and Rad9-depleted NPE (lane 2) were analyzed by immunoblotting with indicated antibodies. Rfc1 was used as a loading control. **(B)** Singly-primed circular ssDNA (lanes 3–12), primed linear ssDNA (lanes 13–22), or linear dsDNA (lanes 23–32) was incubated in NPE shown in (A), and Chk1 phosphorylation was analyzed by immunoblotting with phosphorylated-Chk1 (S344) and Chk1 antibodies. Rfc1 was used as a loading control. The top panel represents a superimposed image (magenta, pChk1; green, Chk1). (*) indicates the heavy chain of rabbit IgG leaked out from IgG beads used for immunodepletion. **(C)** Relative intensities of pChk1 signals were quantified and normalized to that of the 10-min time-point of the mock-treated samples and plotted into a graph. Filled circles represent individual data from two independent experiments, including the one shown in (B). **(D)** Immunodepletion of Topbp1 with a rescue experiment. 0.2 µL each of mock-treated (lane 1) and Topbp1-depleted NPE supplemented with either buffer (lane 2) or 150 nM of recombinant Topbp1 (lane 3) were analyzed by immunoblotting with indicated antibodies. Orc2 was used as a loading control. **(E)** Singly-primed circular ssDNA (lanes 4–15) or linear dsDNA (lanes 16–27) was incubated in NPE shown in (D), and Chk1 phosphorylation was analyzed by immunoblotting. (*) IgG. Note that the levels of leaked IgG vary between mock and ΔTopbp1 conditions depending on the antiserum used for depletion.

The DNA pull-down substrate was prepared as follows: A 3-kb fragment was cut out from pR3k-LacO4 by BsaI and purified through phenol/chloroform extraction, ethanol precipitation, and size exclusion chromatography using the S-400 HR spin column (Cytiva, #27514001). A hairpin oligonucleotide adaptor carrying a Cy5 fluorophore and a biotin moiety (tt2791, Supplementary Table S1) was ligated to one terminus of the fragment, the other terminus was cleaved off with PstI to produce a 4-nt 3′-overhang, and then the 3-kb fragment was separated from oligonucleotides by a Sepharose CL-4B open-top column (Cytiva, #17015001). To append long 3′-ssDNA overhangs, 20 nM of purified fragments were incubated with 0.4 units/µL TdT and 0.1 mM dATP at 37°C for 1 h.

### ATR activation and DSB end resection assays in NPE

NPE was supplemented with 2 mM adenosine triphosphate (ATP), 20 mM phosphocreatine (PC), and 5 μg/mL creatine phosphokinase (CPK) and pre-incubated at 22°C for 5 min. DNA substrates were incubated in NPE at 20 ng/µL at 22°C, and the reaction was stopped at the time points indicated in the figures. To monitor Chk1 phosphorylation, 0.5 µL of the reaction mixture was stopped with 9.5 µL of SDS sample buffer (50 mM Tris-HCl pH 6.8, 0.1 M dithiothreitol, 2% sodium dodecyl sulfate [SDS], 0.05% bromophenol blue, and 10% glycerol), and to monitor DSB end resection, 0.5 µL of the reaction mixture was stopped with 100 μL of stop buffer (1% SDS and 20 mM EDTA). Protein samples, typically 0.2 μL NPE, were separated by SDS polyacrylamide gel electrophoresis (SDS-PAGE), transferred onto polyvinylidene fluoride membranes, and probed with antibodies indicated in the figures. DNA was purified through the treatment with 50 μg/mL Proteinase K at 37°C for 2 h, phenol/chloroform extraction, and ethanol precipitation, and resuspended in TE buffer containing 10 μg/mL RNase A at 1.25 ng/μL. DNA samples were digested with SacI and S1 nuclease (Takara Bio, #2410) and analyzed by agarose gel electrophoresis followed by staining with SYBR Gold nucleic acid gel stain (Thermo Fisher Scientific, #S11494). Fluorescent signals were collected using the Amersham Typhoon scanner 5 system (Cytiva) and processed using ImageJ software (National Institute of Health, MD, USA).

### DNA pull-down assay

Immobilization of linear DNA substrates was carried out essentially as described previously [55], with some modifications. Briefly, 250 ng of Cy5-labeled and biotinylated DNA was incubated with 1 μg of streptavidin (SA, New England Biolabs, #N7021) in 25 μL of binding buffer (10 mM Tris-HCl, pH 7.4, 1 mM EDTA, 1 M NaCl, 0.1% Triton X-100) at 4°C overnight to form DNA-SA complexes, and 100 ng of the DNA-SA complexes were incubated with 1 μL of biotin-sepharose beads in 20 µL of binding buffer at room temperature for 3 h. DNA beads were washed four times with Egg Lysis Buffer salts (ELB-salts: 10 mM Hepes-KOH pH 7.7, 2.5 mM MgCl_2_, 50 mM KCl) and incubated in NPE containing 180 nM GST-S-LacI at the concentration of 20 ng/µL with respect to immobilized DNA at 22°C. At appropriate time points, 1 µL of the reaction mixture was stopped with SDS sample buffer to monitor Chk1 phosphorylation, and 3.5 µL of the mixture was quickly diluted with 200 µL of ELB-salts containing 0.2% Triton X-100, overlayed onto 200 µL of ELB-salts containing 500 mM sucrose, and centrifuged at 12,700 ×*g* for 1 min at 4°C in a TMS-21 swinging bucket rotor (Tomy Seiko, Tokyo, Japan, #0621650333) to monitor DSB end resection and protein binding. The beads were washed once with 200 µL of ELB-salts and resuspended in 14 µL of stop buffer. A 5-µL aliquot was treated with 50 µg/mL proteinase K in stop buffer containing 3 mM biotin to disrupt SA-DNA interaction, and DNA was purified by phenol/chloroform extraction followed by ethanol precipitation, resuspended in TE buffer containing 10 µg/mL RNase A at the concentration of 2.3 ng/µL, digested with Nb.BbvCI, and analyzed by alkaline agarose gel electrophoresis followed by staining with SYBR Gold nucleic acid gel stain. DNA-bound proteins were extracted by SDS sample buffer from the remaining aliquot of the bead mixture and analyzed by SDS-PAGE, followed by immunoblotting.

### Protein expression and purification

Purification of the *Xenopus* 9–1–1 complex was performed as follows: Recombinant proteins were expressed by co-infecting Sf9 insect cells with baculoviruses expressing FLAG-TEV-Rad9, Hus1, and Rad1 for 48 h at 28°C in Sf-900 II SFM (Thermo Fisher Scientific, #0621650333) supplemented with 2% (v/v) fetal bovine serum (FBS). Cells were harvested, washed with phosphate-buffered saline (PBS), and snap-frozen in liquid nitrogen. Cells were suspended in buffer H (25 mM HEPES-NaOH pH 7.8, 1 mM EDTA, 10% glycerol, 0.01% Nonidet P-40) containing 100 μM phenylmethylsulfonyl fluoride (PMSF), 1x cOmplete EDTA-free (Roche, Basel, Switzerland, #5056489001), and 0.1 M NaCl, and the lysates were centrifuged at 100,000 *×g* for 30 min in a Beckman 70.1Ti rotor (Beckman Coulter, CA, USA, #342184). The supernatant was loaded onto a DEAE Sepharose Fast Flow column (Cytiva, #17070901) and eluted with buffer H containing 0.4 M NaCl. The elution fraction was loaded onto an anti-FLAG-M2 agarose column (Sigma Aldrich, MA, US, #A2220), and the bound proteins were eluted with buffer H containing 0.1 M NaCl and 100 μg/mL of FLAG peptide (Sigma Aldrich, #F3290). Peak fractions were pooled and loaded onto a MonoQ 5/50GL column (Cytiva, #17516601), and bound proteins were eluted with a 0.1–0.6 M NaCl linear gradient in buffer H. The 9–1–1 complex in the peak fractions was further purified through size exclusion chromatography using a Superdex 200 Increase 10/300 GL column (Cytiva, #28990944) equilibrated with ELB-salts.

Purification of the GST-S-LacI protein was performed as follows: Recombinant protein expression was induced in *E. coli* BL21-CodonPlus (DE3)-RIPL cells transformed with pColdII-GST-S-LacI by acute temperature shift from 37°C to 15°C and the addition of 0.5 mM Isopropyl β-D-1-thiogalactopyranoside (IPTG) in Terrific broth. After incubation for 24 h at 15°C, cells were harvested, treated with 1 mg/mL lysozyme, and sonicated in buffer S (50 mM Na-phosphate pH 8.0, 500 mM NaCl) containing 1% Triton X-100 and 1 mM PMSF, and centrifuged at 100,000 *×g* for 30 min in a Beckman 70.1Ti rotor. Cleared lysates were passed through a DEAE Sepharose column and then loaded onto a Glutathione Sepharose 4B column (Cytiva, #17075604), and the GST-S-LacI protein was eluted with 10 mM reduced glutathione in buffer H containing 0.1 M NaCl. Peak fractions were pooled and loaded on a MonoQ 5/50 GL column, and bound proteins were eluted with a 0.1–0.6 M NaCl linear gradient in buffer H.

Purification of Topbp1 was performed as follows: Recombinant protein expression was induced in *E. coli* BL21-CodonPlus (DE3)-RIPL cells (Agilent, CA, USA, #230280) transformed with the pET28a-based FLAG-Topbp1 expression plasmid by the addition of 0.5 mM IPTG in Terrific broth for 16 h at 25°C. Cells were harvested, treated with 1 mg/mL lysozyme, sonicated in buffer ST (50 mM Tris-HCl pH 8.0, 0.5 M NaCl, 0.1 % Triton X-100, 5 mM 2-mercaptoethanol, 5% glycerol, 1x cOmplete), and centrifuged at 100,000 ×*g* for 30 min in a Beckman 70.1Ti rotor. Cleared lysates were passed through a DEAE Sepharose column and loaded onto an anti-FLAG M2 agarose column, and FLAG-Topbp1 was eluted with buffer H containing 0.5 M NaCl and 100 μg/mL of the FLAG peptide. The peak protein pool was concentrated by Amicon Ultra-4 (Merck, #UFC801024), loaded onto a 15–35% glycerol gradient in 5 mL of buffer H containing 0.1 M NaCl, and centrifuged using an S52ST rotor (Eppendorf Himac Technologies, Ibaraki, Japan) at 276,000 ×*g* for 20 h at 4°C. Proteins were collected from the bottom of the tube as 200-μL aliquots, and the peak fractions were pooled and concentrated by Amicon Ultra-0.5 mL (Merck, #UFC501024). The S1131A mutant of Topbp1 was expressed and purified as described above except using the pET28a-based FLAG-Topbp1 S1131A expression plasmid for bacterial transformation. All the purified proteins were snap-frozen in liquid nitrogen as small aliquots and stored at -80°C.

### Immunological methods

Production and usage of Rfc1, Orc2, and Msh2 antisera were described previously [56]. The S-tag (BETHYL laboratories, TX, USA, #A190-135A), phospho-Chk1 (Ser345) (Cell Signaling Technology, MA, USA, #2341), which recognizes Ser344 phosphorylation of *Xenopus* Chk1, and Chk1 (G-4) (Santa Cruz Biotechnology, CA, USA, #sc8408) antibodies were commercially available. The ATR antiserum used for immunoblotting and the Topbp1 antiserum have been described previously [57, 58]. Rabbit Rad9, Rad1, Rad50, and Nbs1 antibodies were raised against full-length proteins, rabbit ATM antibodies were raised against the 1–669 fragment of ATM, rabbit Dna2 antibodies were raised against the 1-599 fragment of Dna2, rabbit ATR antibodies used for immunodepletion were raised against the 1-647 fragment of ATR, and rabbit Exo1 antibodies were raised against the full-length, nuclease-deficient mutant (D78A, D137A) of Exo1. All the antigen proteins were expressed in *E. coli* as N-terminally His_6_-tagged proteins. Rabbit Mre11and Rpa2 antibodies were raised against peptides NH_2_-CDEEDFDPFKKSGPSRRGRR-COOH, corresponding to residues 693–711 of Mre11 and NH_2_-CEGHIYSTIDDEHYKSTDGD-COOH, corresponding to residues 258–276 of Rpa2, respectively. For immunoblotting, Rad9, Rad1, Rad50, Nbs1, Topbp1, ATM, ATR, Mre11, Rfc1, Orc2, Exo1, Dna2, and Rpa2 antisera were used at 1:5,000, pChk1 (S345) antibodies were used at 1:1,000, and Chk1 antibodies were used at 1:500 dilutions, respectively. Alexa Fluor 647-conjugated Goat anti-rabbit IgG (H+L) antibodies (Jackson ImmunoResearch, PA, USA, #111-605-144) and Alexa Fluor 488-conjugated Goat anti-mouse IgG (H+L) antibodies (Jackson ImmunoResearch, #115-165-146) were used at 1:10,000 dilutions as the secondary antibodies. To evaluate the depletion efficiency, 0.2 μL of NPE was loaded on SDS-PAGE unless otherwise indicated.

For immunodepletion from NPE, 3 vol of an antiserum was bound to 1 vol of recombinant Protein A Sepharose Fast Flow (PAS; Cytiva, #17127902) at 4°C overnight, 0.2 vol of the antibody-coupled PAS beads were incubated with 1 vol of NPE at 4°C for 1 h, and the procedure was repeated twice. For double depletion, 1.5 vol each of the two sera was bound to 1 vol of PAS, 0.3 vol of antibody-coupled PAS beads were incubated with 1 vol of NPE at 4°C for 1 h, and the procedure was repeated twice. In a typical experimental setup, we mixed 18 µL of an antiserum and 6 µL of PAS at 4°C overnight, 2 µL of the antibody-coupled PAS beads were incubated with 10 µL of NPE at 4°C for 1 h, and the procedure was repeated twice.

## RESULTS

### Linear double-stranded DNA activates ATR through a 9–1–1-independent pathway in *Xenopus* egg extracts

To investigate the regulation of DSB response, we took advantage of the nucleoplasmic extract of *Xenopus* eggs (NPE), in which primed circular ssDNA and linear dsDNA have been used to activate ATR [39, 59]. We first tested how different DNA structures affect ATR checkpoint activation pathways in this system. To this end, we immunodepleted the 9–1–1 complex, a DNA structure sensor for the ATR pathway, from NPE, and incubated various DNA substrates in the mock-treated and Rad9-depleted extracts (Figure 1A). As shown previously [59], primed circular ssDNA efficiently induced Chk1 serine 344 (S344) phosphorylation, a readout of ATR activation, in a Rad9-dependent manner (Figure 1B, top panel, see “primed ss circular”). Linear dsDNA also efficiently induced Chk1 S344 phosphorylation. However, Rad9-depletion reduced Chk1 phosphorylation only modestly in this case, suggesting that linear dsDNA activates a 9– 1–1-independent ATR checkpoint pathway (Figures 1B and 1C). A key determinant seems to be DNA termini, since once linearized, the primed ssDNA substrate evoked detectable, although not strong, Chk1 phosphorylation in Rad9-depleted NPE (Figures 1B and 1C, compare “primed ss circular” and “primed ss linear”). In contrast to 9–1–1, both primed ss circular and ds linear substrates required Topbp1 to induce Chk1 S344 phosphorylation (Figures 1D and 1E). These results suggest that linear dsDNA induces a 9–1–1-independent but Topbp1-dependent ATR activation pathway in *Xenopus* egg extracts.

### Redundant roles of 9–1–1 and MRN in ATR activation and DSB end resection

We next set out to identify factors required for the 9–1–1-independent ATR activation pathway induced by linear dsDNA. MRN was an obvious candidate since MRN has been reported to activate ATR in response to DSBs [38, 42] and primed circular ssDNA [39]. However, as MRN is critical for the initiation of DSB end resection, the depletion of MRN would prevent DSB end resection, and as a consequence, compromise ssDNA-mediated ATR activation. To bypass the MRN’s role in the initiation of DSB end resection, we prepared a resection-intermediate-like DNA substrate, which has been shown to bypass the requirement of MRN in the initiation of DSB end resection [47], by appending ssDNA overhangs of several hundred base pairs to the 3′-termini. We then incubated this DNA substrate, which we call “tailed DNA” hereafter, in NPE. The DNA products were then recovered and treated with S1 nuclease to measure the remaining dsDNA length (Figure 2A). As expected, when blunt-ended “tailless” DNA was used as a substrate, Mre11-depletion inhibited DSB end resection and increased the NHEJ products (Figures 2B and 2C, see “tailless”). Chk1 phosphorylation was not observed in Mre11-depleted NPE (Figure 2D). In contrast, tailed DNA induced efficient Chk1 phosphorylation, produced almost no NHEJ products, and received additional end resection even in the absence of Mre11, indicating that 3′-ssDNA overhangs indeed bypassed the initiation of DSB end resection (Figures 2C and 2D, “tailed”). It would be worth noting that we avoided using exonucleases to make 3′-ssDNA overhangs since they are in general processive enzymes whose products tend to be a mixture of differently processed fragments. Instead, we used terminal deoxynucleotidyl transferase (TdT) to extend 3′-ssDNA overhangs since TdT is a distributive enzyme whose products are relatively uniform in length (Supplementary Figure S1).

**Figure 2.**
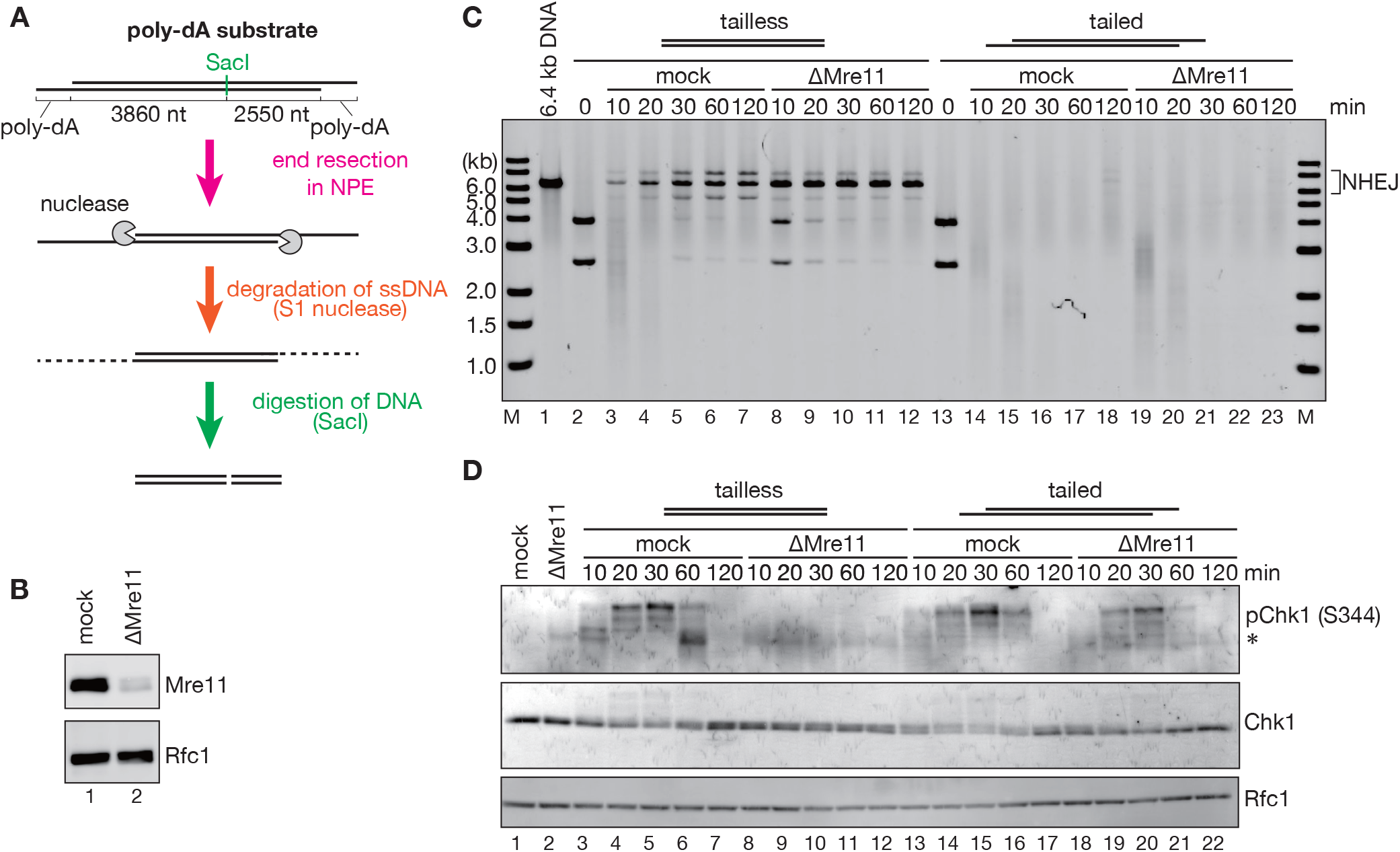
Bypass of the requirement of MRN in the initiation of DSB end resection by 3′-ssDNA overhangs. **(A)** Schematic diagram of the DNA substrate and the DSB end resection assay. The substrate was incubated in NPE, purified, and digested with S1 nuclease to measure the length of the dsDNA portion and with SacI to monitor end resection at two termini separately. **(B)** 0.2 µL each of mock-treated (lane 1) and Mre11-depleted NPE (lane 2) were analyzed by immunoblotting with indicated antibodies. Rfc1 was used as a loading control. **(C)** The tailless substrate (lanes 2–12) or the tailed substrate (lanes 13– 23) was incubated in NPE shown in (B), processed as described in (A), and analyzed by agarose gel electrophoresis. The DNA fragment of approximately 6.4 kb corresponds to the major head-to-tail NHEJ product. **(D)** The tailed substrate was incubated in NPE shown in (B), and Chk1 phosphorylation was analyzed by immunoblotting. (*) IgG.

Using the tailed DNA substrate, we next asked whether MRN backs up ATR activation in the absence of 9–1–1. As seen with tailless linear DNA, tailed DNA induced robust Chk1 phosphorylation in Rad9-depleted NPE (Figures 3A and 3B). Strikingly, when both Mre11 and Rad9 were simultaneously depleted, Chk1 phosphorylation was reduced to a nearly undetectable level (Figures 3B and 3C). Although the level of Chk1 phosphorylation in Rad9/Mre11-doubly-depleted NPE varies between experiments likely due to the technical difficulty of double depletion (for example, see Figure 3F), we consistently observed a significant reduction of Chk1 phosphorylation by Rad9/Mre11 double-depletion. Furthermore, to our surprise, DSB end resection was significantly slowed down in the absence of Mre11 and Rad9 (Figures 3D and 3E). The *Xenopus* 9–1–1 complex purified from insect cells restored Chk1 phosphorylation and the rate of DSB end resection at least partially (Figures 3F, 3G, and Supplementary Figures S2A–C). Likewise, immunodepletion of Nbs1 phenocopied that of Mre11 with respect to both DSB end resection and Chk1 phosphorylation, suggesting that the observed phenotype reflects the MRN function (Figures 3H, 3I, and Supplementary Figures S2D–F). These results strongly suggest that MRN and 9–1–1 redundantly promote both ATR activation and long-range DSB end resection in response to DSB substrates.

**Figure 3.**
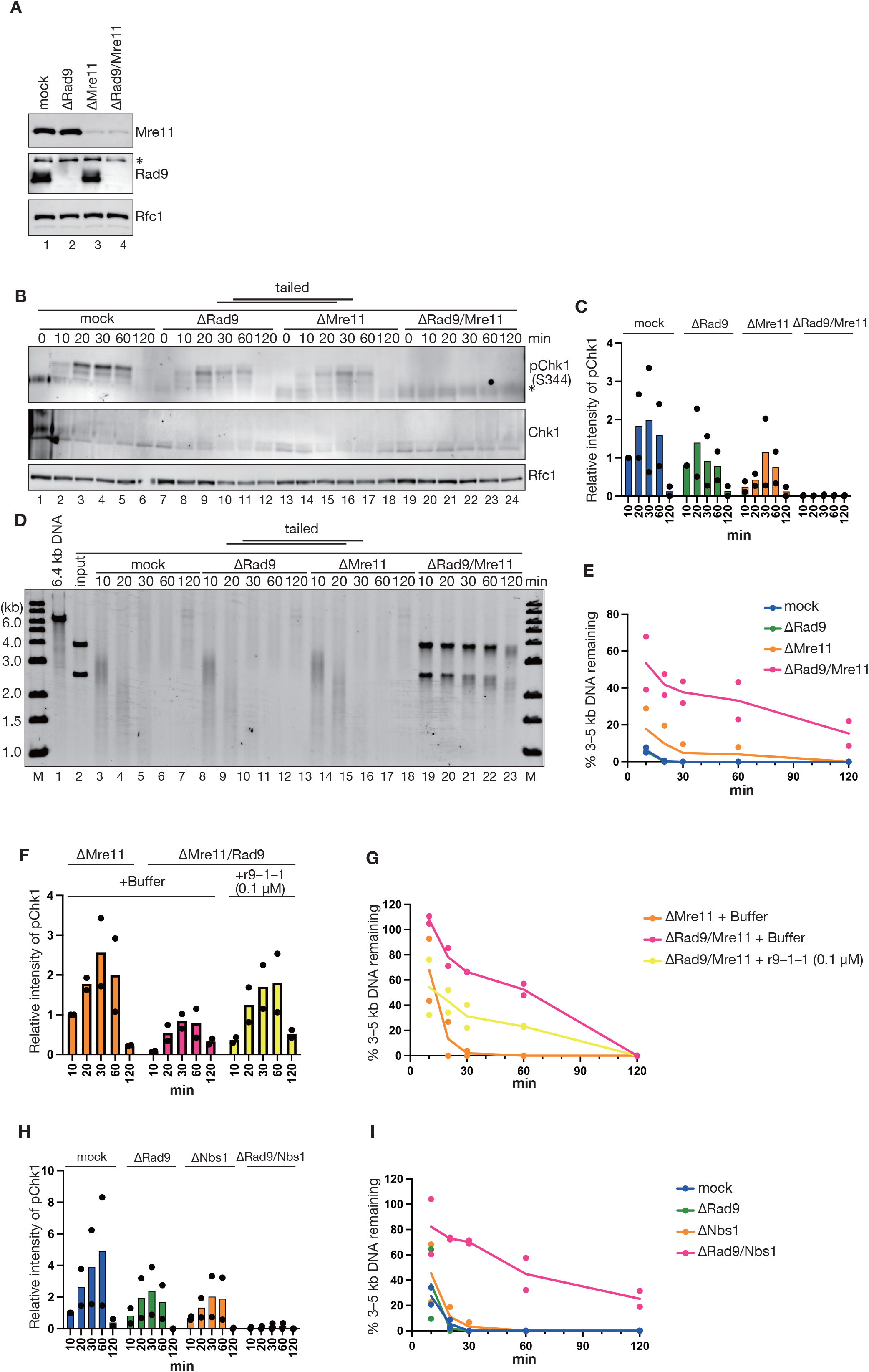
Redundant roles of 9–1–1 and MRN in ATR activation and DSB end resection. **(A)** 0.2 µL each of mock-treated (lane 1), Rad9-depleted (lane 2), Mre11-depleted (lane 3), and Rad9/Mre11-doubly-depleted NPE (lane 4) were analyzed by immunoblotting with indicated antibodies. Rfc1 was used as a loading control. (*) Cross-reacting band. **(B)** The tailed substrate was incubated in NPE shown in (A), and Chk1 phosphorylation was analyzed by immunoblotting. (*) IgG. **(C)** Quantification of Chk1 phosphorylation for (B). Relative intensities of pChk1 signals were quantified, normalized to that of the 10-min time-point of the mock-treated samples, and plotted into a graph. Filled circles represent individual data from two independent experiments, including the one shown in (B). **(D)** The tailed substrate was incubated in NPE shown in (A) and analyzed as described in Figure 2A. **(E)** Quantification of long-range end resection for (D). Relative intensities of the DNA signals within the 3-to 5-kb range were quantified, normalized to the intensity of the input DNA, and plotted into a graph. Filled circles represent individual data from two independent experiments, including the one shown in (D). **(F)** Relative intensities of pChk1 signals in response to tailed DNA in Mre11- and Rad9/Mre11-depleted NPE with a rescue experiment of recombinant 9–1–1, processed as described in (C). See Supplementary Figure S2B for a representative image. **(G)** Quantification of long-range end resection in Mre11- and Rad9/Mre11-depleted NPE with a rescue experiment of recombinant 9–1–1, processed as described in (E). See Supplementary Figure S2C for a representative image. **(H)** Relative intensities of pChk1 signals in response to tailed DNA in mock-, Rad9-, Nbs1-, and Rad9/Nbs1-depleted NPE, processed as described in (C). See Supplementary Figure S2E for a representative image. **(I)** Quantification of long-range end resection in mock-, Rad9-, Nbs1- and Rad9/Nbs1-depleted NPE, processed as described in (E). See Supplementary Figure S2F for a representative image.

### ATM is an essential component of the MRN-dependent ATR activation pathway

MRN is a well-known ATM activator. To test whether MRN activates the ATR checkpoint pathway through ATM, we next depleted ATM from NPE in combination with Rad9 and Mre11 (Figure 4A). The depletion of ATM modestly reduced, and ATM/Rad9-double-depletion strongly inhibited Chk1 phosphorylation (Figures 4A–C). In contrast, ATM/Mre11-double-depletion did not further reduce Chk1 phosphorylation compared to single-depletion of either one (Figures 4B and 4C, compare ΔMre11, ΔATM, and ΔMre11/ΔATM). From these data, we infer that MRN and ATM participate in the same ATR activation pathway.

**Figure 4.**
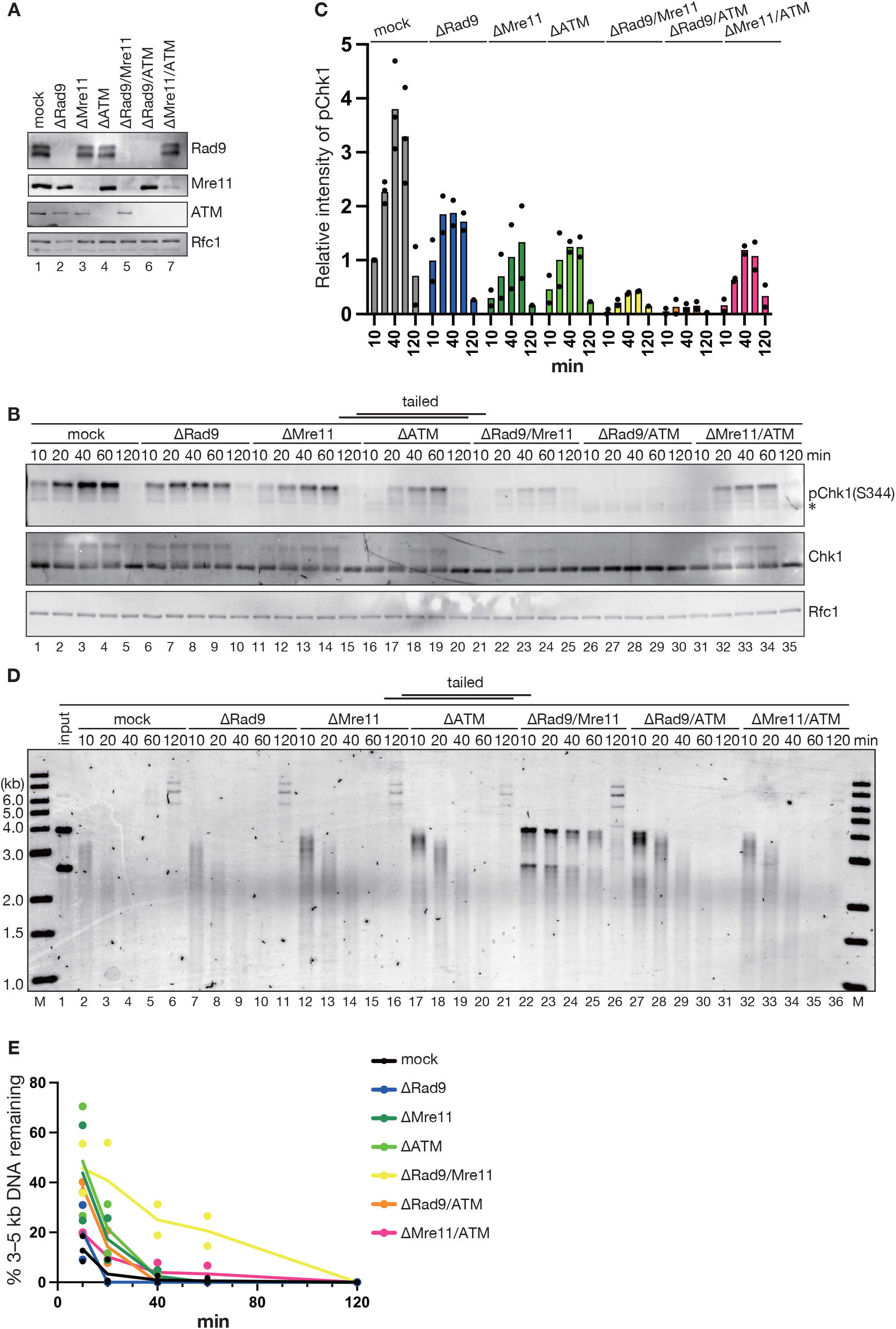
ATM is an essential component of the MRN-dependent ATR activation pathway. **(A)** 0.2 µL each of mock-treated (lane 1), Rad9-depleted (lane 2), Mre11-depleted (lane 3), ATM-depleted (lane 4), Rad9/Mre11-doubly-depleted (lane 5), Rad9/ATM-doubly-depleted (lane 6), and Mre11/ATM-doubly-depleted NPE (lane 7) were analyzed by immunoblotting with indicated antibodies. Rfc1 was used as a loading control. **(B)** The tailed substrate was incubated in NPE shown in (A), and Chk1 phosphorylation was analyzed by immunoblotting. (*) IgG. **(C)** Quantification of Chk1 phosphorylation for (B). Data were processed and presented as described in Figure 1C. **(D)** The tailed substrate was incubated in NPE shown in (A) and analyzed as described in Figure 2A. **(E)** Quantification of long-range end resection for (D). Data were processed and presented as described in Figure 3E.

Interestingly, although Rad9-depletion strongly attenuated long-range DSB end resection when combined with Mre11-depletion, it showed only a minimal effect on the rate of end resection when combined with ATM-depletion (Figures 4D and 4E). These data argue that although MRN regulates both ATR activation and DSB end resection, ATM supports only the checkpoint function of MRN.

### Checkpoint sensors stimulate long-range DSB end resection largely independently of checkpoint kinases

To further address how long-range DSB end resection is regulated by checkpoint kinases, we next depleted ATM and ATR from NPE. Neither ATR-nor ATM-depletion detectably reduced the rate of DSB end resection, although the simultaneous depletion of both moderately attenuated DSB end resection (Supplementary Figures S3). These results suggest that 9–1–1 and MRN stimulate long-range end resection in a pathway that is largely independent of the ATM/ATR checkpoint kinases. While it is possible that two checkpoint kinases have redundant roles in regulating long-range end resection, this point needs further verification including functional rescue of the phenotype with recombinant proteins.

### 9–1–1 and MRN stimulate the Dna2-dependent DSB end-resection pathway through distinct mechanisms

To gain further insight into how 9–1–1 and MRN promote ATR activation and DSB end resection, we set up a DNA pull-down assay (Figure 5A). A 3′-ssDNA overhang was appended to one terminus of a 3-kb linear DNA by TdT, and the other terminus was capped with a hairpin structure to prevent end resection from this terminus. We also placed a Cy5 fluorophore for detection, a biotin moiety in the hairpin loop to immobilize DNA, a nicking endonuclease site to separate the top and bottom strands, and four tandem *lac* operator (*lacO*) repeats for the normalization purpose. The DNA was immobilized on biotin-conjugated Sepharose beads through streptavidin and incubated in NPE with recombinant GST- and S-tagged LacI used as a loading control. On an alkaline agarose gel electrophoresis, we confirmed that the Cy5-carrying, 5′-terminated top strand was gradually degraded while the 3′-terminated bottom strand remained intact in NPE (Figure 5B, see “mock”). Resection of the top strand was attenuated in NPE depleted of both Rad9 and Mre11 (Figure 5B). Chk1 phosphorylation was reduced partially by depleting either Rad9 or Mre11 and drastically by depleting both from NPE (Figure 5C). Therefore, the immobilized DNA substrate recapitulated all the essential phenotypes observed with the free substrate.

**Figure 5.**
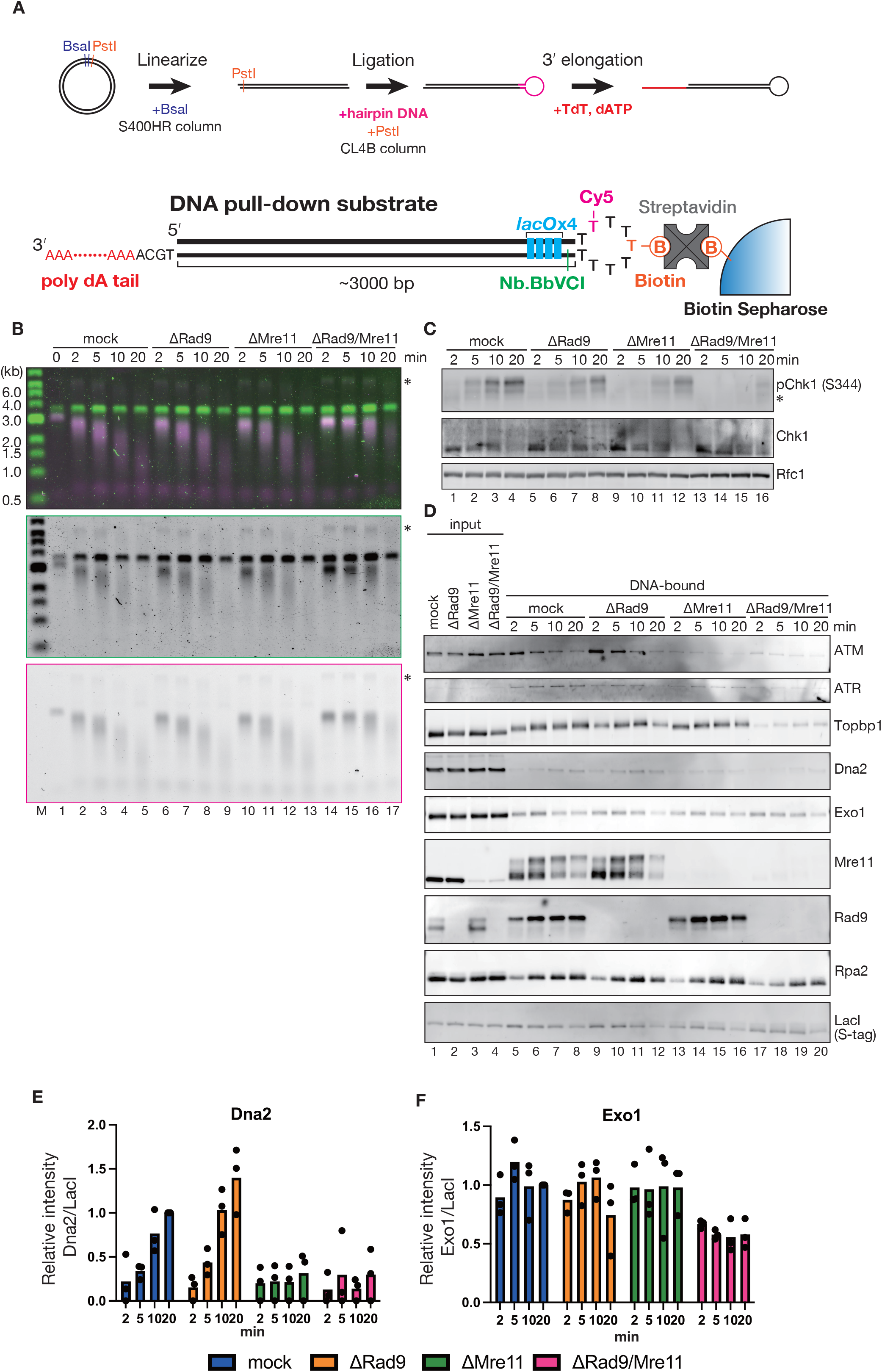
Recruitment of checkpoint and end-resection factors on tailed DNA. **(A)** A diagram of the substrate for the DNA pull-down assay. A 3-kb plasmid was linearized with BsaI, and a hairpin adaptor carrying Cy5- and biotin-modified bases was ligated to one terminus. Another terminus was cleaved with PstI to make a 4-nt 3′-overhang, and a poly-dA tail was appended to this terminus with TdT. The resulting DNA was immobilized on biotin-Sepharose through tetrameric streptavidin as a linker. **(B)** The substrate shown in (A) was immobilized on beads, incubated with GST-S-LacI in NPE depleted of mock, Rad9, Mre11, or both, purified, treated with the Nb.BbvCI nickase to separate two strands, and analyzed by alkaline agarose gel electrophoresis. The DNA was visualized by SYBR-Gold (middle), and the resected DNA strand was monitored by Cy5 fluorescence (bottom). The top panel represents a superimposed image (magenta, Cy5; green, SYBR-Gold). (*) indicates substrates escaped from Nb.BbvCI digestion. **(C)** Immunoblotting for Chk1 phosphorylation. (*) IgG. **(D)** The immobilized substrate was recovered and bound proteins were analyzed by immunoblotting with indicated antibodies. LacI was used as a loading control. **(E, F)** Relative intensities of DNA-bound Dna2 (E) or Exo1 signals (F) were quantified, normalized to that of the 20-min time-point of the mock-treated samples, and plotted into a graph. Filled circles represent individual data from three independent experiments, including the one shown in (D).

Using this system, we sought the mechanism of how 9–1–1 and MRN stimulate end resection. Loading of Rpa2 gradually increased during the time course, likely reflecting the exposure of ssDNA (Figure 5D, “mock”). Long-range DSB end resection is mediated by Dna2 and Exo1, among which the former is the major resection nuclease in *Xenopus* egg extracts [60]. We confirmed that Dna2-depletion strongly attenuates long-range end resection (Supplementary Figure S4). Furthermore, we found that both MRN and 9–1–1 redundantly stimulate the Dna2-dependent pathway, since in the absence of Exo1, where only the Dna2-dependent pathway remains active, the depletion of Rad9 or Mre11 did not inhibit DSB end resection to the level seen in Mre11/Rad9-doubly-depleted extracts (Supplementary Figure S4). Interestingly, however, the contribution of MRN and 9–1–1 to Dna2 loading was different at the molecular level; Rad9-depletion did not detectably affect the loading of Dna2 nor Exo1, whereas Mre11-depletion significantly reduced the loading of Dna2 onto immobilized DNA (Figures 5D–F). These results argue that 9–1–1 and MRN stimulate the resection nucleases through distinct mechanisms; MRN enhances the loading of Dna2 onto DNA, whereas 9–1–1 stimulates its activity mostly independently of its loading.

### The assembly of checkpoint complexes on immobilized DNA carrying a 3′-ssDNA overhang

The bead-binding assay also provided information on how 9–1–1 and MRN activate the ATR checkpoint pathway. Firstly, the loading of Mre11 was not affected in the absence of Rad9, and Rad9 loading was largely retained in the absence of Mre11, indicating that they are loaded onto tailed DNA mostly through independent mechanisms (Figure 5D). Secondly, the loading of Topbp1 was reduced drastically by depleting both Rad9 and Mre11, strongly suggesting that 9–1–1 and MRN redundantly promote Topbp1-loading. Thirdly, we observed clear ATM loading onto tailed DNA, and this loading depended on Mre11 (Figure 5D, compare “mock” and “ΔMre11”). Interestingly, ATM loading peaked at an earlier time point and quickly went down in the later time points not only on DNA carrying a short 4-nt overhang but also on DNA carrying a long 3′-ssDNA overhang, implying that the length of the 3′-ssDNA overhang is not a critical determinant for the ATM loading and unloading kinetics (Supplementary Figure S5). Overall, these data suggest that 9–1–1 and MRN independently facilitate the loading of Topbp1, while MRN also transiently recruits ATM even in the presence of a long 3′-ssDNA overhang.

### The mechanism of ATR activation through the MRN–ATM–Topbp1 pathway

Our data suggest that the MRN-dependent ATR activation pathway functions through ATM and Topbp1, consistent with previous reports [38, 42, 44]. It has been found that ATM-dependent phosphorylation of a serine residue (serine 1131 in *Xenopus laevis*) in AAD of Topbp1 is critical for Chk1 phosphorylation induced by poly(dA)_70_–(dT)_70_ oligonucleotides [44]. To investigate the role of serine 1131 (S1131) phosphorylation in our system, we prepared wild-type (Topbp1^WT^) and the phosphorylation-deficient mutant of Topbp1 (Topbp1^S1131A^). As expected, Topbp1 was essential for Chk1 phosphorylation in response to both tailless and tailed DNA fragments and poly(dA)_70_– (dT)_70_ synthetic oligonucleotides and was not essential for DSB end resection (Figures 6A, 6B, and Supplementary Figures S6A and S6B). Consistent with previous reports, Topbp1^WT^ but not Topbp1^S1131A^ restored Chk1 phosphorylation in response to poly(dA)_70_–(dT)_70_, confirming that phosphorylation of S1131 is critical for ATR activation induced by this substrate (Figure 6B). Interestingly, Topbp1^S1131A^ partially restored Chk1 phosphorylation in response to the tailed DNA substrate (Figure 6B, “tailed”), likely because the tailed substrates activate ATR through 9–1–1 and MRN, only the latter of which relies on ATM (Figure 4). To separate these two pathways, we next depleted Topbp1 in combination with Rad9 and Mre11. In the absence of Mre11, where the 9–1– 1 pathway should dominate, both Topbp1^WT^ and Topbp1^S1131A^ restored Chk1 phosphorylation in response to tailed DNA, suggesting that S1131 phosphorylation is not essential for ATR activation in the 9–1–1 pathway (Supplementary Figures S6C and S6D). Strikingly, Topbp1^S1131A^ also restored Chk1 phosphorylation, albeit partially, in the extract depleted of Rad9, where the MRN/ATM-dependent pathway should dominate (Figures 6C and 6D). These results suggest that, during long-range end resection, MRN activates ATR through two subpathways; one dependent on Topbp1 S1131 phosphorylation and one independent of it.

**Figure 6.**
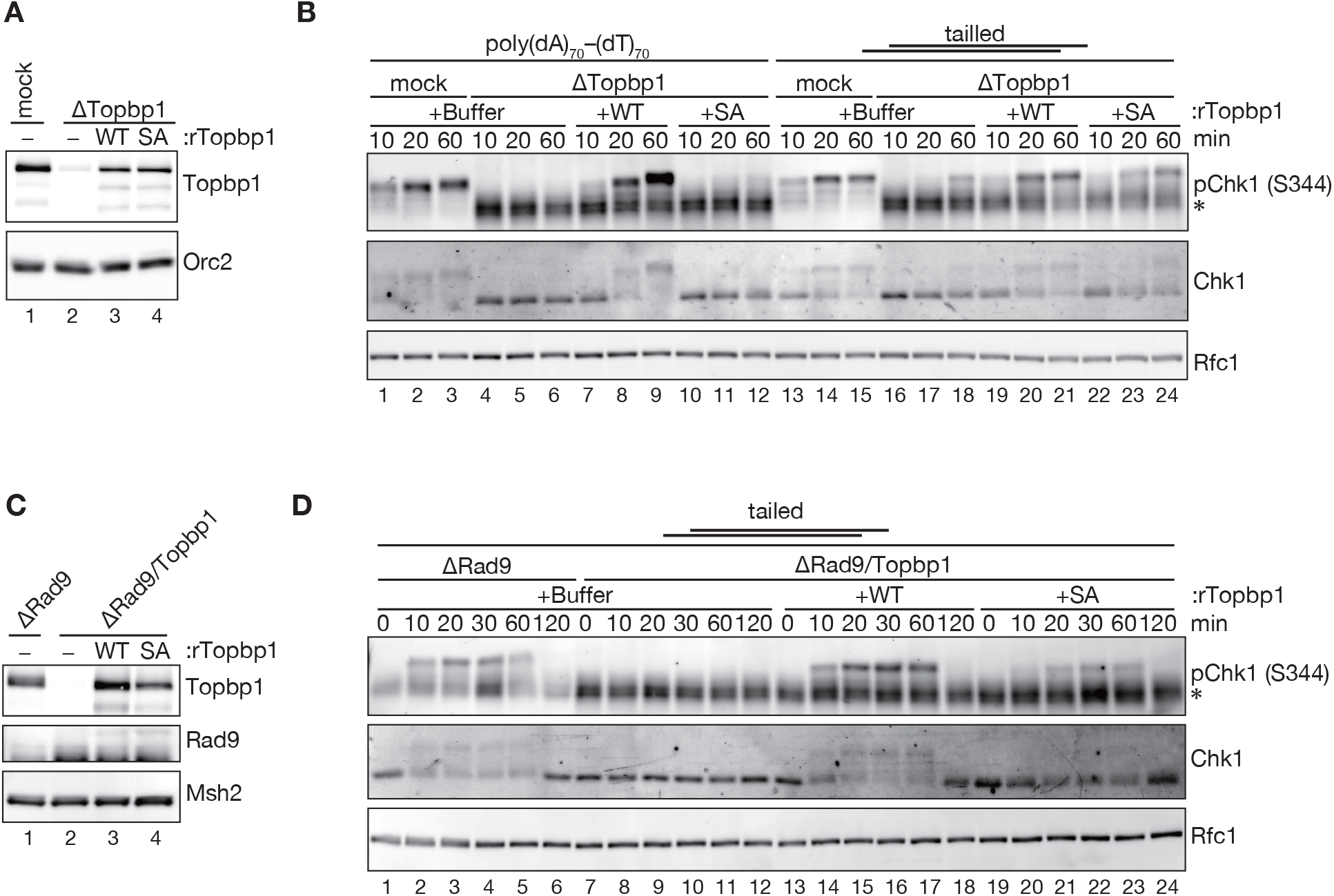
Requirement of the serine 1131 phosphorylation site in Topbp1 in MRN- and 9–1–1-dependent ATR activation subpathways. **(A)** Immunodepletion of Topbp1 with a rescue experiment. 0.2 µL each of mock-treated (lane 1) and Topbp1-depleted NPE supplemented with either buffer (lane 2) or 150 nM of recombinant Topbp1 (WT: wild-type, lane 3; SA: S1131A, lane 4) were analyzed by immunoblotting with indicated antibodies. Orc2 was used as a loading control. **(B)** Poly(dA)_70_–(dT)_70_ oligonucleotides or the tailed substrate were incubated in NPE shown in (A), and Chk1 phosphorylation was analyzed by immunoblotting. (*) IgG. **(C)** Immunodepletion of Topbp1 and Rad9 with a Topbp1 rescue experiment. 0.2 µL each of Rad9-depleted (lane 1) and Rad9/Topbp1-doubly-depleted NPE (lanes 2–4) supplemented with either buffer (lane 2) or 150 nM of recombinant Topbp1 (WT: wild-type, lane 3; SA: S1131A, lane 4) were analyzed by immunoblotting with indicated antibodies. Msh2 was used as a loading control. **(D)** The tailed substrate was incubated in NPE shown in (C), and Chk1 phosphorylation was analyzed by immunoblotting. (*) IgG.

### Topbp1-mediated regulation of checkpoint complex assembly

Since Topbp1 plays a central role in the activation of ATR, we asked how it affects the assembly of checkpoint complexes using the DNA pull-down assay described in Figure 5. Interestingly, the depletion of Topbp1 had a significant impact on the loading and dissociation of checkpoint proteins on DNA with a long 3′-ssDNA overhang; the loading of Rad9 and ATR was greatly reduced, ATM was accumulated on DNA, and phosphorylation of Mre11 was significantly impaired (Figure 7A and Supplementary Figure S7). These results suggest that Topbp1 stabilizes a multiprotein complex including ATR and 9–1–1 and promotes Mre11 phosphorylation but stimulates the unloading of ATM.

**Figure 7.**
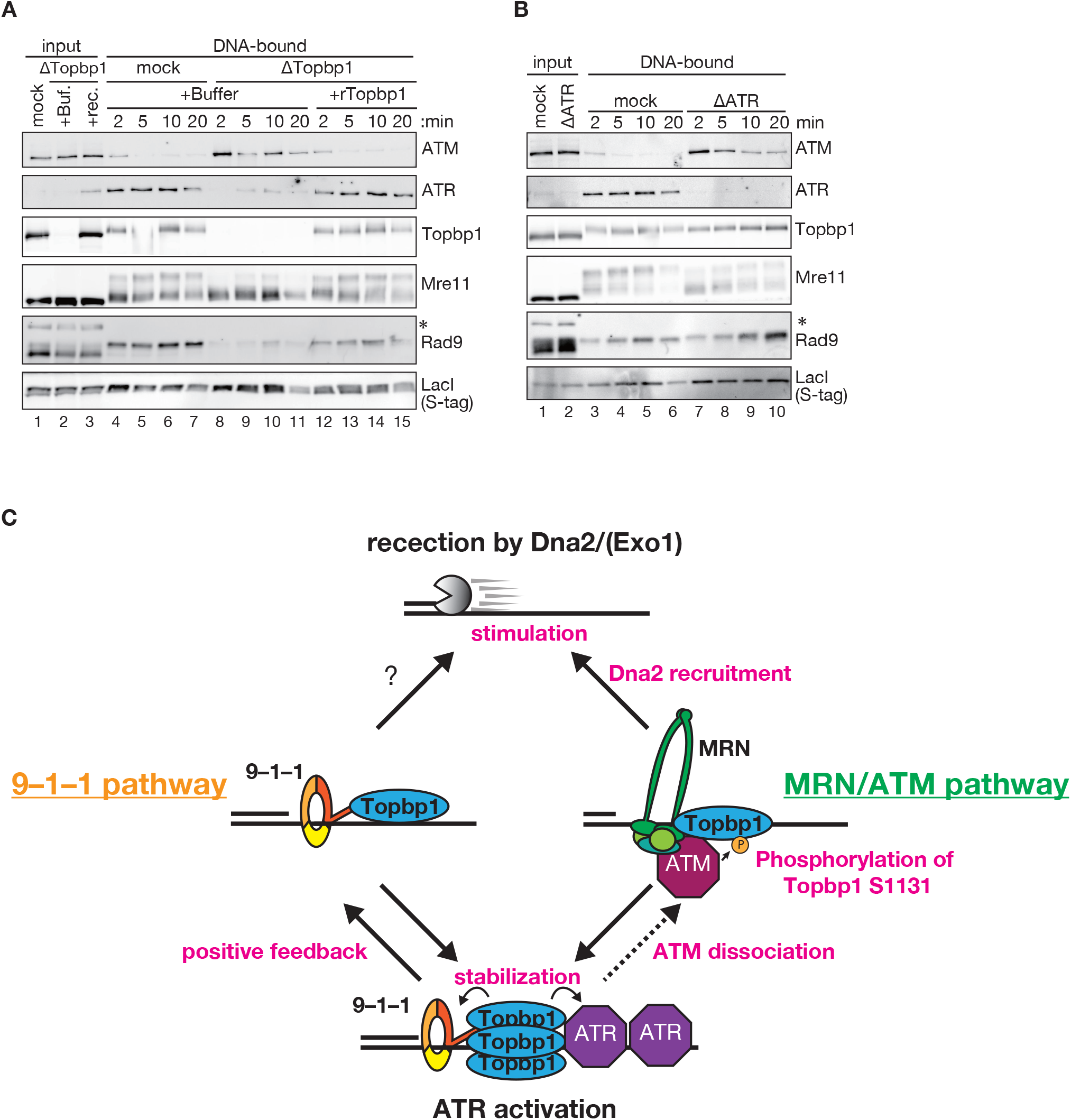
Requirement of Topbp1 and ATR in the assembly of checkpoint complexes on tailed DNA. **(A)** The tailed substrate shown in Figure 5A was immobilized on beads, incubated with GST-S-LacI in NPE depleted of mock (lanes 1 and 4–7) or Topbp1 (lanes 2, 3, and 8–15) supplemented either buffer (lanes 1, 2 and 4–11) or 150 nM recombinant Topbp1 (lanes 3 and 12–15), and recovered. Input (lanes 1–3) and DNA-bound samples (lanes 4–15) were analyzed by immunoblotting with indicated antibodies. LacI was used as a loading control. **(B)** The tailed substrate shown in Figure 5A was immobilized on beads, incubated with GST-S-LacI in NPE depleted of mock (lanes 1 and 3–6) or ATR (lanes 2 and 7–10), and recovered. Input (lanes 1 and 2) and DNA-bound samples (lanes 3–10) were analyzed by immunoblotting with indicated antibodies. LacI was used as a loading control. **(C)** A model for the 9–1–1- and MRN-dependent ATR activation and DSB end resection pathways. See text for details.

To dissect multiple roles of Topbp1 in the regulation of checkpoint complexes, we next depleted ATR from NPE. In contrast to Topbp1-depletion, the loading of Rad9, as well as that of Topbp1, was constant in the absence of ATR (Figure 7B). However, the depletion of ATR impaired both Mre11 phosphorylation and ATM unloading. These data argue that Topbp1 promotes Mre11 phosphorylation and ATM ejection through the activation of ATR, whereas it stabilizes the association of 9–1–1 and ATR on DNA during long-range end resection independently of the function of ATR.

## DISCUSSION

Both 9–1–1 and MRN play multiple roles in DNA damage response, including ATR activation and DSB end resection. Thus, MRN activates ATR by recruiting Topbp1 and mediating ATM-dependent Topbp1 phosphorylation, 9–1–1 recruits Topbp1 to activate ATR, and both MRN and 9–1–1 stimulate the resection nucleases [13, 34-36, 38-40, 47, 49, 50, 61]. Since these mechanisms have been separately studied in humans, *Xenopus*, and yeasts, a large part of the relationship between these reactions has been uncertain. In this study, using DNA substrates that mimic extensively resected DSBs, we showed that these distinct mechanisms operate simultaneously and redundantly during long-range resection in *Xenopus* egg extracts. By establishing a bead-based DNA loading assay, we further provided molecular mechanisms of how 9–1–1 and MRN promote ATR checkpoint activation and DSB end resection. Our data pinpoint Topbp1 as the critical organizer of both 9–1–1- and MRN-dependent ATR activation pathways and Dna2 as the major end-resection nuclease activated by the two pathways.

Although it has been known that both ATM- and ATR-dependent checkpoint pathways regulate DSB end resection [2], our data clearly indicate that 9–1–1 and MRN facilitate the end-processing reaction independently of checkpoint activation. This is in good agreement with previous biochemical data demonstrating that 9–1–1 and MRN enhance the activity of Exo1 and Dna2–RecQ helicase [13, 15, 45-47, 49, 62]. Since bacterial homologs of Rad50 and Mre11 form a rigid clamp upon DNA binding [63], MRN may serve as the loading platform for the Dna2-RecQ helicase complex, as previously suggested [13, 15, 46]. Likewise, 9–1–1 is a ring-shaped clamp that is loaded onto 5′-P/T junctions by Rad17-RLC [20-24], and a recent strand-specific ChIP analysis showed that 9–1–1 dynamically relocates on dsDNA with ongoing end resection and remains close to 5′-P/T junctions [64]. 9–1–1 may stimulate Dna2 by tethering it around 5′-P/T junctions, in a manner such that the PCNA replication clamp holds DNA polymerases. Interestingly, our data argue that MRN is important for the recruitment or stability of the majority of Dna2, whereas 9–1–1 is largely dispensable for the loading of Dna2 (Figure 5). The interpretation should be careful since this may reflect the number of molecules of the two sensors on DNA. Although MRN can form multimers on DNA [65], 9–1–1 is the checkpoint clamp that may be present as a monomer at a 5′-P/T junction, leaving it possible that our assay underestimates the contribution of 9–1–1 to Dna2 loading. In contrast to Dna2, the loading of Exo1 was reduced by the depletion of Rad9 and Mre11 only partially (Figure 5F). Since DSB end resection depends mostly on the Dna2 pathway in this system, we were not able to test rigorously the contribution of 9–1–1 and MRN to Exo1 recruitment and stimulation, which was observed previously [13, 45, 62].

Our DNA pull-down assay identified Topbp1 as the critical organizer of the two checkpoint-activation pathways. In good agreement with previous findings [34-36, 38-40, 61], our data fit well with the scenario that MRN recruits ATM and Topbp1, and 9–1– 1 recruits Topbp1, to activate ATR (Figures 5 and 7). Interestingly, in the absence of Topbp1, we found that the loading of 9–1–1 and ATR are significantly compromised. Since 9–1–1 is loaded onto 5′-P/T junctions by Rad17-RLC and ATR–ATRIP is recruited onto ssDNA through RPA, Topbp1 is unlikely required for their initial loading steps. Therefore, Topbp1 likely stabilizes 9–1–1 and ATR after their initial recruitment, possibly through multivalent protein-protein interactions and forming protein condensates [30]. In contrast to the strong effect on 9–1–1 and ATR, Topbp1-depletion did not destabilize MRN and ATM on DNA. It instead stabilized the loading of ATM, which was otherwise transient but clearly seen on the tailed DNA substrate that carries no DSB termini (Figure 7). Topbp1-depletion also reduced Mre11 phosphorylation (Figure 7A and Supplementary Figure S7). Since MRN directly interacts with ATM [3, 8], ATR-dependent phosphorylation of Mre11 possibly turns off the ATM signaling pathway by dissociating ATM from MRN. Consistently, Mre11 phosphorylation has been reported to be inhibitory for downstream events, including ATM activation and resection by Exo1 [66-69]. The fact that the switching from the ATM to the ATR checkpoint pathway is critical for proper DSB response warrants further verification of this hypothesis.

Figure 7C summarizes our model. After the initial short-range end resection by the MRN complex, two redundant mechanisms involving the 9–1–1 and MRN checkpoint sensors activate the ATR checkpoint signaling pathway and the DSB long-range end processing machinery. We assume that the activation of the MRN pathway is specific to DNA substrates that have termini, involves the recruitment of ATM, and partially requires phosphorylation of Topbp1 on serine 1131. Both 9–1–1 and MRN redundantly promote Topbp1 loading. Topbp1, in turn, stabilizes 9–1–1 and ATR and promotes the dissociation of ATM from DNA through the activation of ATR. Two pathways also stimulate Dna2 loading and activation mostly independently of the ATM/ATR signaling pathway. We propose that these two redundant pathways ensure the robust and efficient activation of the ATR signaling cascade in combination with DSB end processing.

## DATA AVAILABILITY

The data underlying this article are available in the article and in its online supplementary material. Unprocessed raw data and DNA sequences underlying this article will be shared on reasonable request to the corresponding authors.

## SUPPLEMENTARY DATA

Supplementary Data are available at NAR online.

## AUTHOR CONTRIBUTIONS

Kensuke Tatsukawa: Conceptualization, Formal analysis, Investigation, Methodology, Writing—original draft. Reihi Sakamoto: Investigation, Methodology. Yoshitaka Kawasoe: Methodology. Yumiko Kubota: Methodology. Toshiki Tsurimoto: Conceptualization, Funding acquisition. Tatsuro S. Takahashi: Conceptualization, Funding acquisition, Supervision, Methodology, Writing—review & editing. Eiji Ohashi: Conceptualization, Funding acquisition, Supervision, Writing—review & editing.

## ACKNOWLEDGEMENTS

We thank Dr. Bunsho Shiotani for critical reading of the manuscript, Dr. Asako Furukohri for sharing materials and technical information, and Mr. Eiichiro Kanatsu for the expression and purification of Exo1 used for raising the antibodies. We also thank members of the Takahashi lab for helpful discussions.

## FUNDING

This work was supported by JSPS KAKENHI grants [22H04697, 20K21399, and 20H03186 to T.S.T], [16H04743 to T.T], [19K06613 to E.O], and [22K15036 to Y. Kawasoe.], and a grant-in-aid for JSPS fellows [21J14041 to K.T]. This work was also supported by the Toyota Riken Scholar program [to E.O., T.S.T., and Y. Kawasoe], the Uehara Memorial Foundation research grant [to Y. Kawasoe], and the R3 Young Researchers Support Project, Faculty of Science, Kyushu University Grant [21-07 to Y. Kawasoe].

## CONFLICT OF INTEREST

The authors declare no competing interest.

## SUPPLEMENTARY FIGURE LEGENDS

**Supplementary Figure S1.**
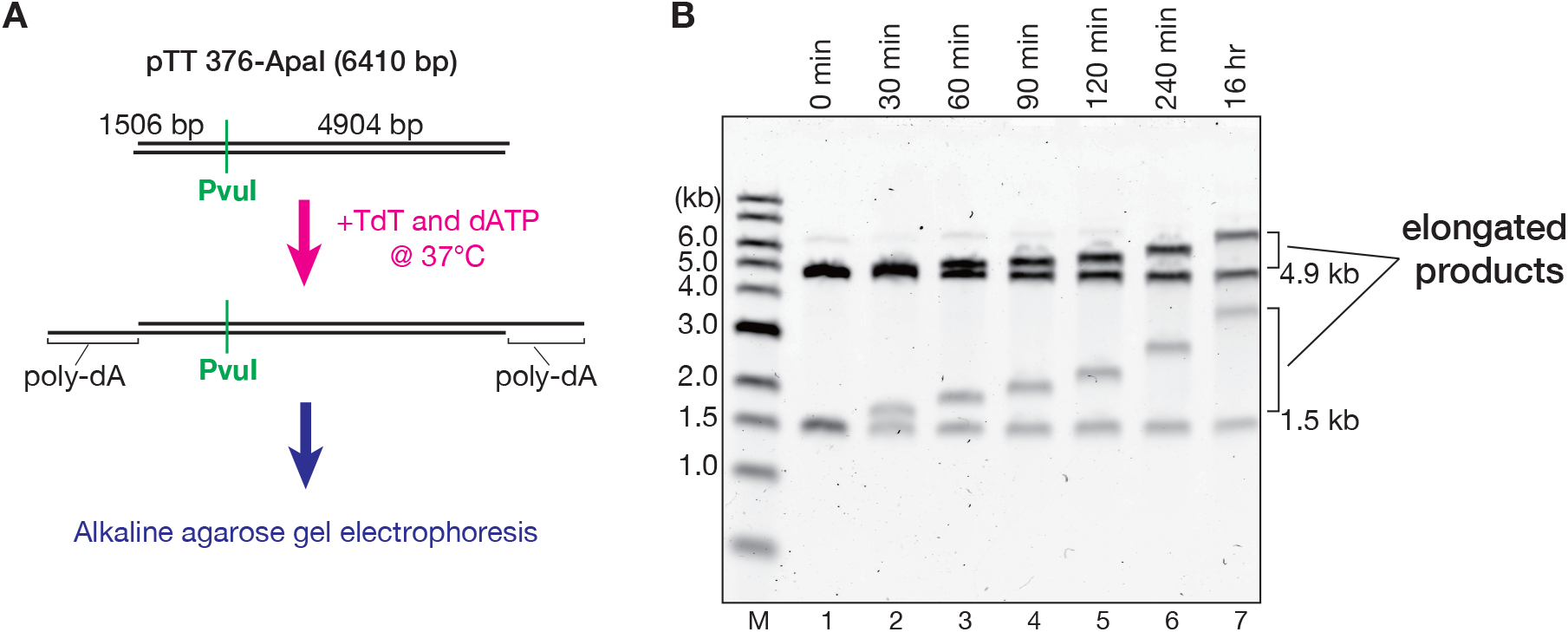
Preparation of tailed DNA substrates. **(A)** A schematic diagram of the preparation of DNA substrates carrying 3′-ssDNA overhangs. Blunt-ended linear dsDNA was incubated with terminal deoxynucleotidyl transferase (TdT) and dATP at 37°C. The length of the 3′-ssDNA overhangs was estimated by alkaline agarose gel electrophoresis after digestion with PvuI. **(B)** The reaction products analyzed by alkaline agarose gel electrophoresis. Relatively uniform extension of a specific strand was observed over time.

**Supplementary Figure S2.**
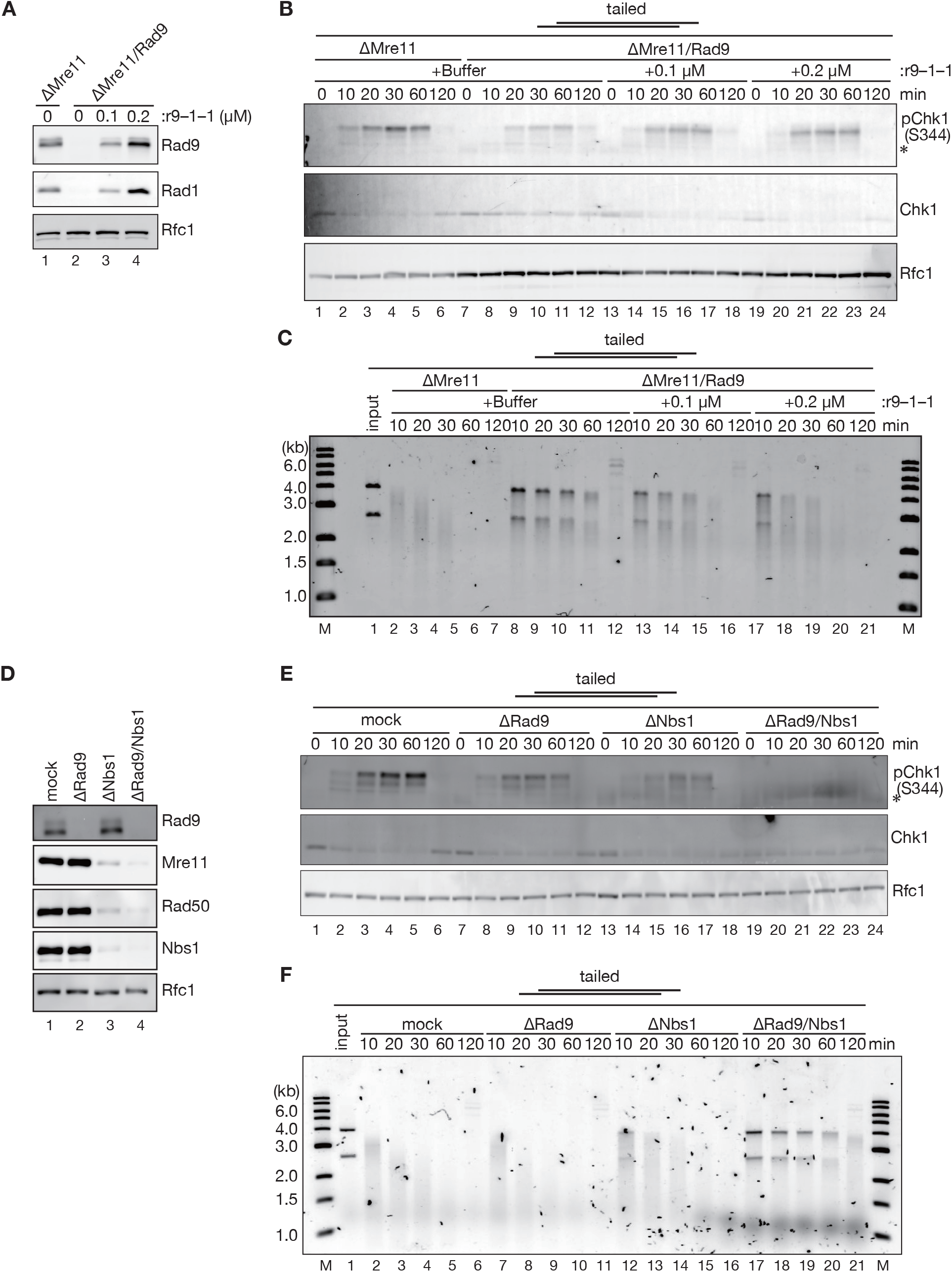
Redundant roles of 9–1–1 and MRN in ATR activation and DSB end resection. **(A)** Immunodepletion of Rad9 and Mre11 with a 9–1–1 rescue experiment. 0.2 µL each of Mre11-depleted (lane 1) and Mre11/Rad9-doubly-depleted NPE (lanes 2–4) supplemented with either buffer (lanes 1 and 2) or recombinant 9–1–1 (lane 3, 100 nM; lane 4, 200 nM) were separated by SDS-PAGE and probed with indicated antibodies. Rfc1 was used as a loading control. **(B)** The tailed substrate was incubated in NPE shown in (A) and Chk1 phosphorylation was analyzed by immunoblotting. (*) IgG. **(C)** The tailed substrate was incubated in NPE and analyzed as described in Figure 2A. **(D–F)** The effect of Rad9/Nbs1 double-depletion on Chk1 phosphorylation and long-range DSB end resection. The experiments shown in Figures 3A–C were repeated using the Nbs1 antiserum instead of the Mre11 antiserum. Immunoblotting for the depletion efficiency using 0.2 μL each of NPE (D), immunoblotting for Chk1 phosphorylation (E), and agarose gel electrophoresis for end resection of the tailed substrate (F) were presented. Nbs1-depletion, in combination with Rad9-depletion, strongly suppressed Chk1 phosphorylation and DSB end resection as seen for Rad9/Mre11-double depletion.

**Supplementary Figure S3.**
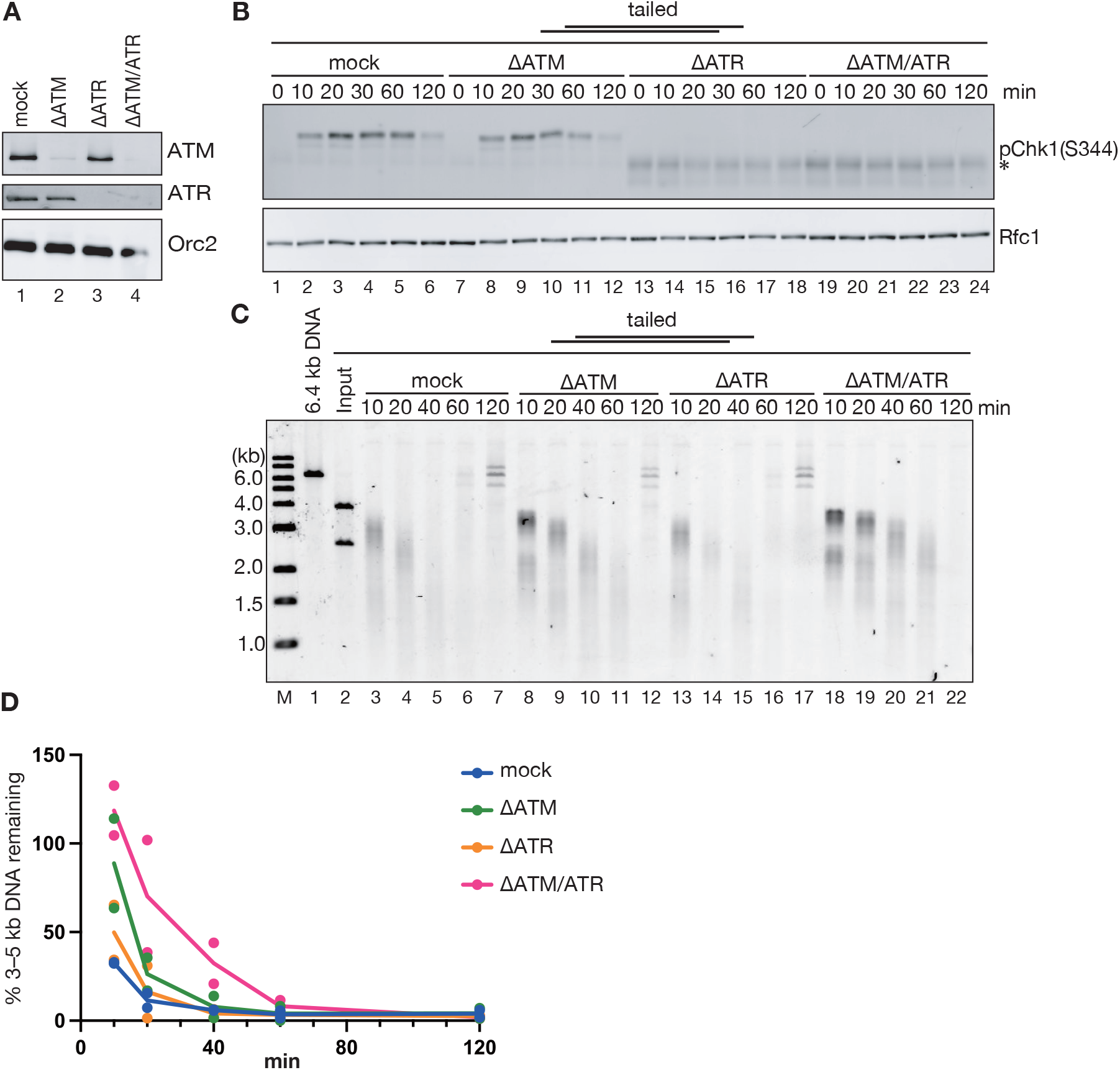
The effect of depletion of checkpoint kinases on long-range DSB end resection. **(A)** 0.2 µL each of mock-treated (lane 1), ATM-depleted (lane 2), ATR-depleted (lane 3), and ATM/ATR-doubly-depleted NPE (lane 4) were analyzed by immunoblotting with indicated antibodies. Orc2 was used as a loading control. **(B)** The tailed substrate was incubated in NPE shown in (A) and Chk1 phosphorylation was analyzed by immunoblotting. (*) IgG. **(C)** The tailed substrate was incubated in NPE shown in (A) and analyzed as described in Figure 2A. **(D)** A graph representing relative intensities of 3–5 kb DNA fragments for (C), normalized to input DNA, with raw data from two replicates shown by filled circles, including the one shown in (C). Immunodepletion of ATM or ATR did not show a strong impact on the rate of long-range end resection, although double-depletion of both ATM and ATR slowed down the resection rate.

**Supplementary Figure S4.**
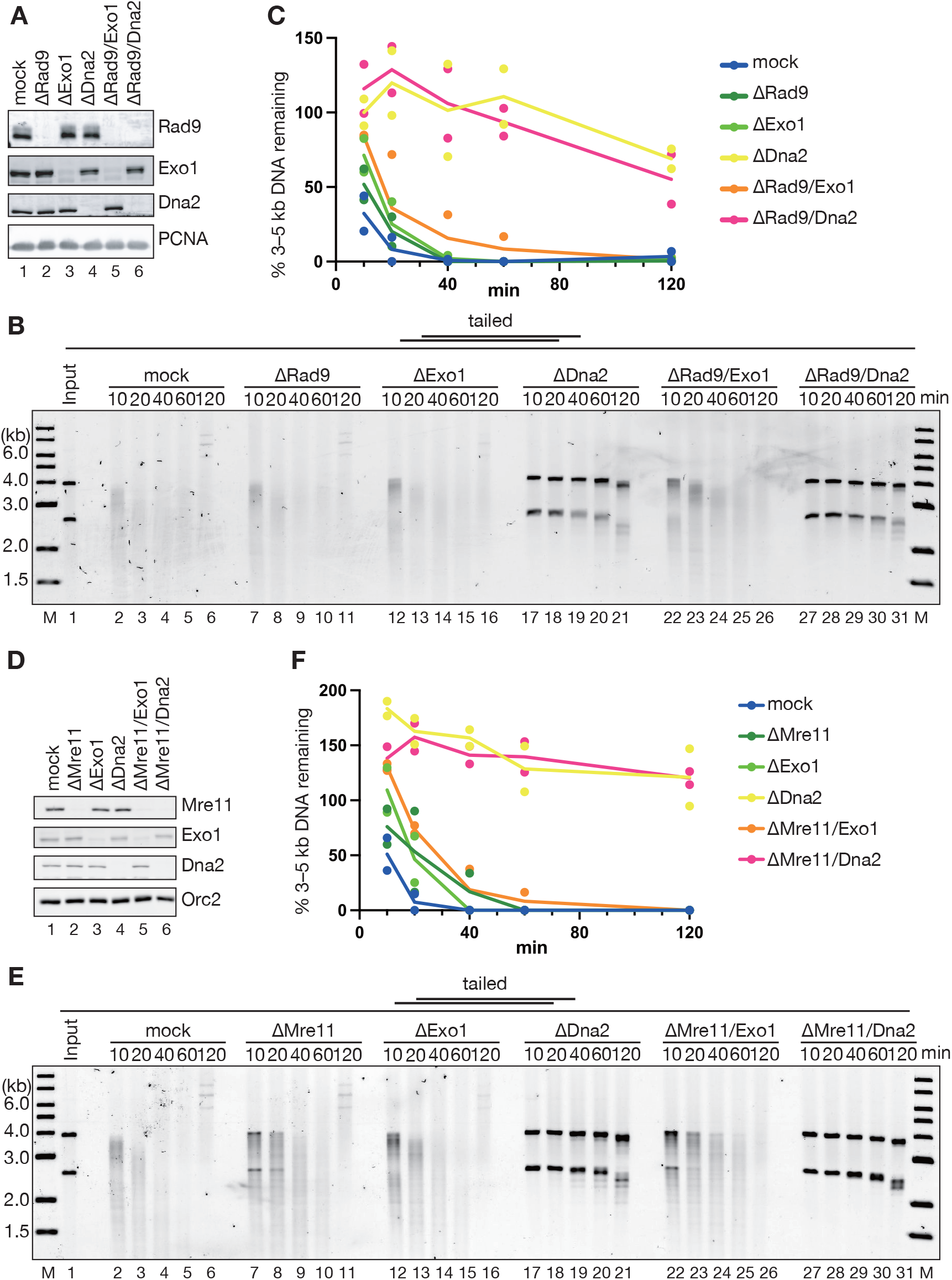
9–1–1 and MRN stimulate the Dna2-dependent DSB resection pathway. **(A–C)** The effect of Rad9/Exo1- and Rad9/Dna2-double-depletion on DSB end resection. The tailed DNA substrate was incubated in NPE depleted of Rad9, Exo1, and Dna2, either alone or in combination, and the DNA products were analyzed by agarose gel electrophoresis. Immunoblotting for the depletion efficiency using 0.2 μL each of NPE (A) and an agarose gel image for end resection (B) were presented. (C) represents a graph showing intensities of 3–5 kb DNA fragments relative to input DNA, with raw data from two replicates shown by filled circles, including the one shown in (B). Dna2-depletion significantly delayed the rate of DSB end resection, while Exo1-depletion had a minor impact on it, suggesting that Dna2 is the major end-resection nuclease in *Xenopus* egg extracts. **(D–F)** The experiments shown in (A–C) were repeated using the Mre11 antiserum instead of the Rad9 antiserum. Immunoblotting for the depletion efficiency (D), an agarose gel image for end resection (E), and a graph showing intensities of 3–5 kb DNA fragments relative to input DNA (F) were presented. These data collectively suggest that both 9–1–1 and MRN stimulate the Dna2-dependent end resection pathway.

**Supplementary Figure S5.**
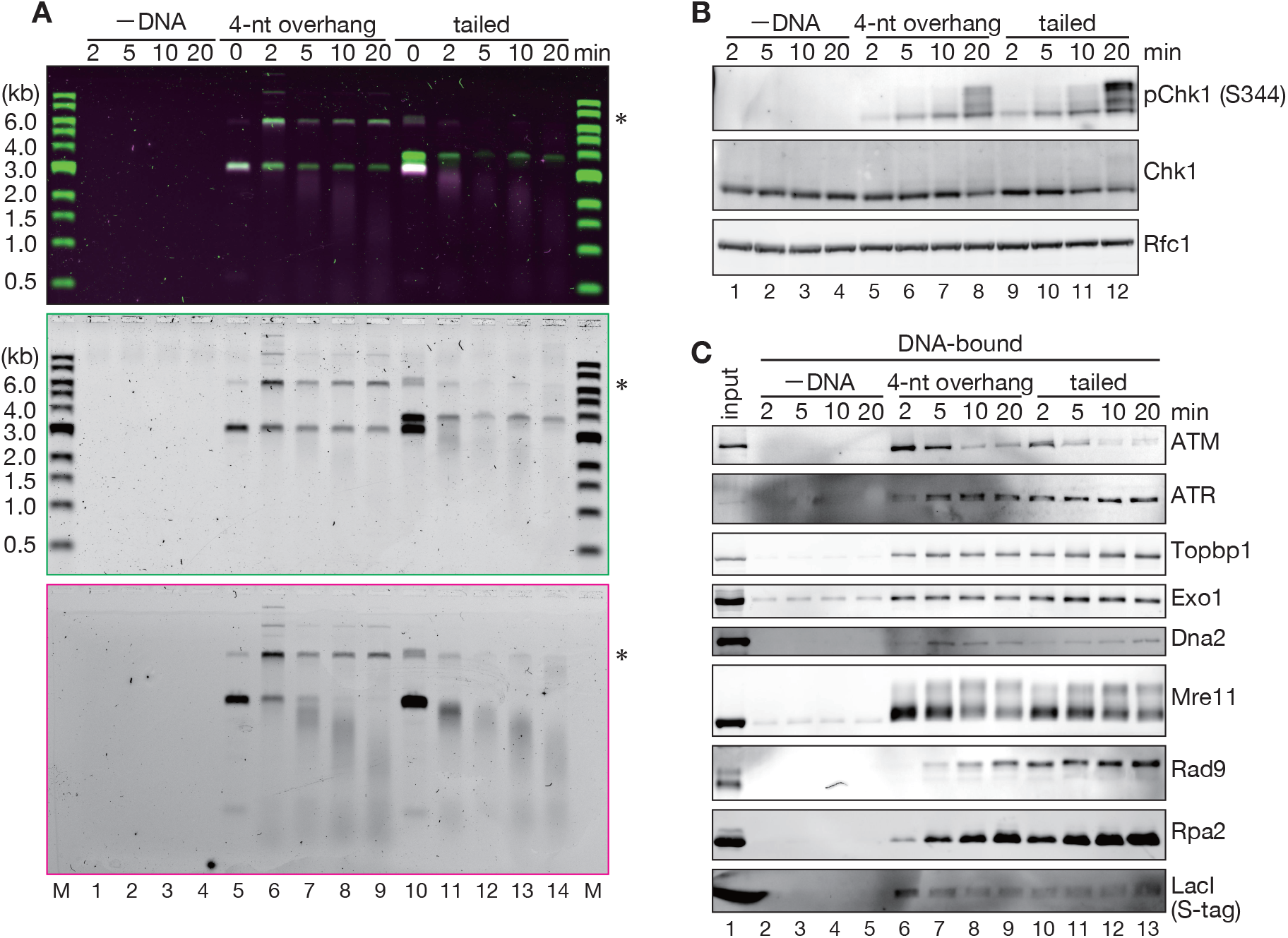
DNA pull-down assay with DNA substrates carrying different lengths of 3′-ssDNA overhangs. **(A)** The DNA substrates carrying 4-nt and long 3′-ssDNA overhangs shown in Figure 5A were immobilized on Sepharose beads, incubated in NPE in the presence of GST-S-LacI, and analyzed as described in Figure 5B. DNA was visualized by SYBR-Gold (middle), and the 5′-terminated strand was detected with Cy5 fluorescence (bottom). The top panel represents a superimposed image (magenta, Cy5; green, SYBR-Gold). The 6-kb fragments indicated by asterisks are either NHEJ products or substrates escaped from Nb.BbvCI digestion. Long 3′-ssDNA overhangs suppressed NHEJ and accelerated the initiation of DSB end resection (compare lanes 6 and 11). **(B)** Chk1 phosphorylation in the reaction mixture shown in (A). **(C)** The DNA pull-down assay. Long 3′-ssDNA overhangs stimulated the loading of Rad9 and Rpa2, whereas reduced the loading of ATM. Notwithstanding the presence of 3’-ssDNA overhangs that prevent NHEJ, the tailed substrate supported the efficient loading of Mre11, and while reduced, ATM was transiently loaded onto tailed DNA at early time points.

**Supplementary Figure S6.**
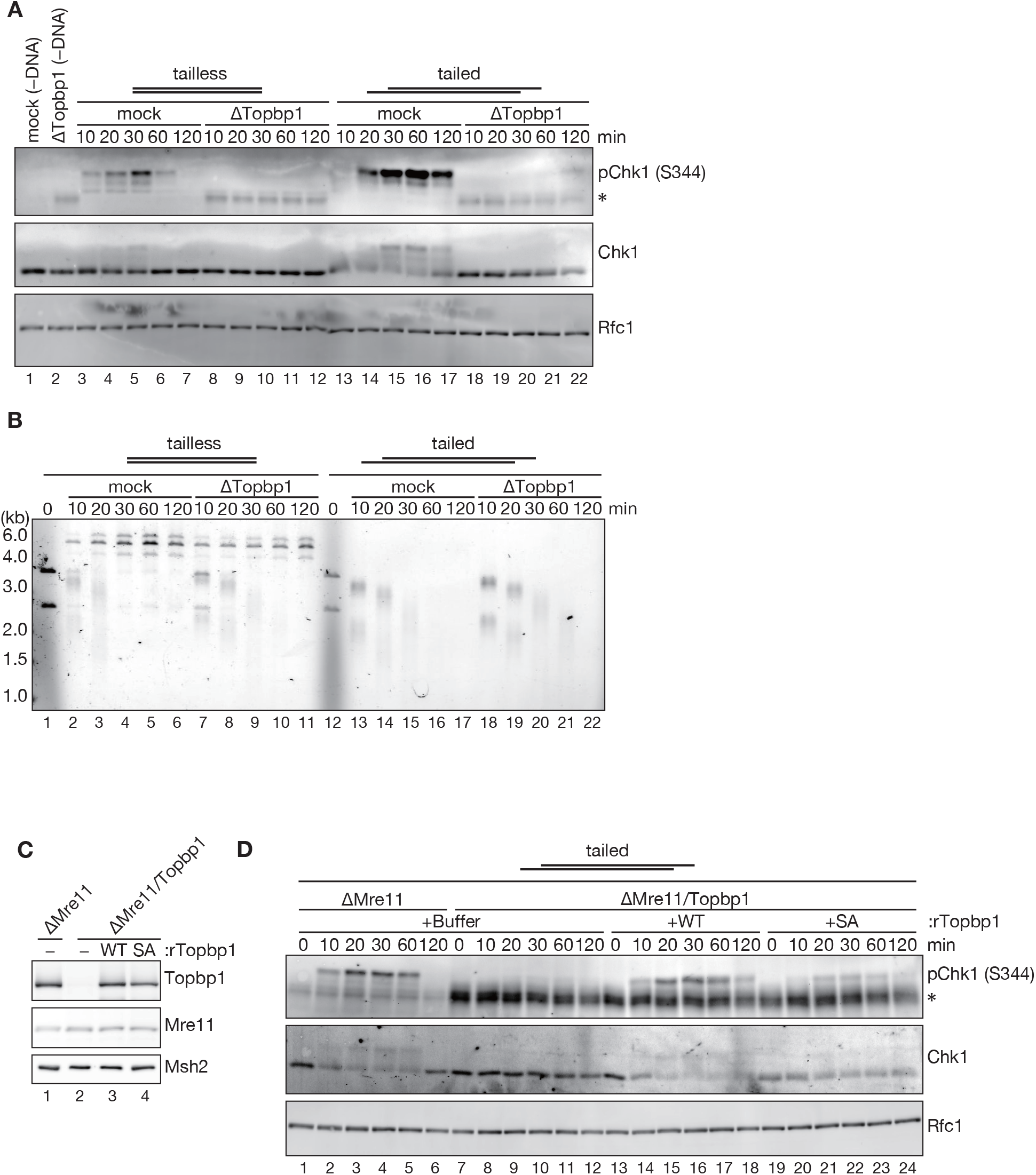
Requirement of Topbp1 in ATR-checkpoint activation and long-range DSB end resection. **(A)** The tailless or the tailed substrates was incubated in mock-treated or Topbp1-depleted NPE and Chk1 phosphorylation was analyzed by immunoblotting. **(B)** The DSB end resection assay. **(C)** Immunodepletion of Topbp1 and Mre11 with a Topbp1 rescue experiment. Immunoblots of NPE samples are shown. Msh2 was used as a loading control. **(D)** Chk1 phosphorylation in response to the tailed substrate.

**Supplementary Figure S7.**
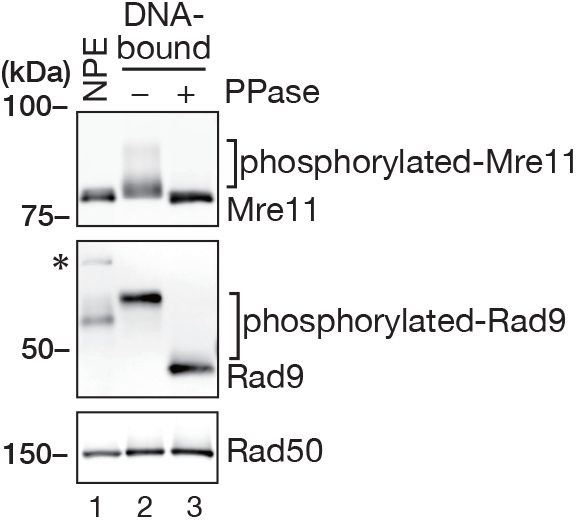
Phosphorylation of Mre11 and Rad9. DNA-bound proteins were recovered from NPE using immobilized DNA carrying 3′-ssDNA overhangs as described in Figure 5A at 10 min after incubation, treated with either buffer (lane 2) or lambda protein phosphatase (lane 3), eluted with SDS-PAGE sample buffer, analyzed by SDS-PAGE alongside 0.2 μl NPE (lane 1), and probed with Mre11, Rad9, and Rad50 antibodies. Both Mre11 and Rad9 showed phosphatase-sensitive retardation of gel electrophoretic mobility, suggesting that the slow-migrating bands are their phosphorylated forms.

## Notes

### Competing Interest Statement

The authors have declared no competing interest.

